# Adaptive robustness through incoherent signaling mechanisms in a regenerative brain

**DOI:** 10.1101/2023.01.20.523817

**Authors:** Samuel R. Bray, Livia S. Wyss, Chew Chai, Maria E. Lozada, Bo Wang

**Author notes:** these authors contributed equally to this work (Samuel R. Bray, Livia S. Wyss).

## Abstract

Animal behavior emerges from collective dynamics of interconnected neurons, making it vulnerable to connectome damage. Paradoxically, many organisms maintain significant behavioral output after large-scale neural injury. Molecular underpinnings of this extreme robustness remain largely unknown. Here, we develop a quantitative behavioral analysis pipeline to measure previously uncharacterized long-lasting latent memory states in planarian flatworms during whole-brain regeneration. By combining >20,000 animal trials with neural population dynamic modeling, we show that long-range volumetric peptidergic signals allow the planarian to rapidly reestablish latent states and restore coarse behavior after large structural perturbations to the nervous system, while small-molecule neuromodulators gradually refine the precision. The different time and length scales of neuropeptide and small-molecule transmission generate incoherent patterns of neural activity which competitively regulate behavior and memory. Controlling behavior through opposing communication mechanisms creates a more robust system than either alone and may serve as a generic approach to construct robust neural networks.

## Introduction

Given its high interconnected complexity, the nervous system is expected to be vulnerable to major neuronal losses such as injuries, stroke, and degeneration^1,2^. However, many animals are capable of regenerating large sections of their nervous system after severe injury while maintaining high levels of motor function and sensitivity to various stimuli^3–7^. The extreme robustness of their nervous system allows them to sense and escape from harmful environmental cues such as predators and UV irradiation even during the process of regrowing a head. Neural robustness is generally thought to be built in the topology of synaptic connectome using redundant links to remove nodes of high centrality and reduce dependency on any given neuron^1,2,8–10^. Examples include distributed nerve nets of cnidarians^3,4^ and duplicated neural circuits in segmented animals such as annelids and insects^5,6^. However, the duplication of network components may be limited by the high metabolic cost of neural maintenance^11^.

Here, we present evidence for an alternative strategy for neural robustness: in addition to synaptic connections, volumetrically transmitted long-range signals could increase effective connectivity of the network without adding new structures to the system (**Figure 1A**). Volume transmission is common in the nervous system and occurs at multiple scales. Besides transmitting across synapses, small molecules such as monoamines and acetylcholine can leak out of the synaptic cleft and function as neuromodulators. However, due to rapid reuptake and extracellular degradation, their diffusion is limited to fast timescales (~100 ms) and short distances (~µm), thereby targeting immediately adjacent neurons^12–14^. In contrast, neuropeptides can be secreted throughout the entire neuronal body and diffuse for up to minutes over hundreds of microns, transmitting their signal to potentially large numbers of neurons with matching receptors^15–17^. The large length scale of neuropeptide communication and its independence from synaptic connections may reduce sensitivity to disruptions like missing neurons, axons, or connections.

**Figure 1:**
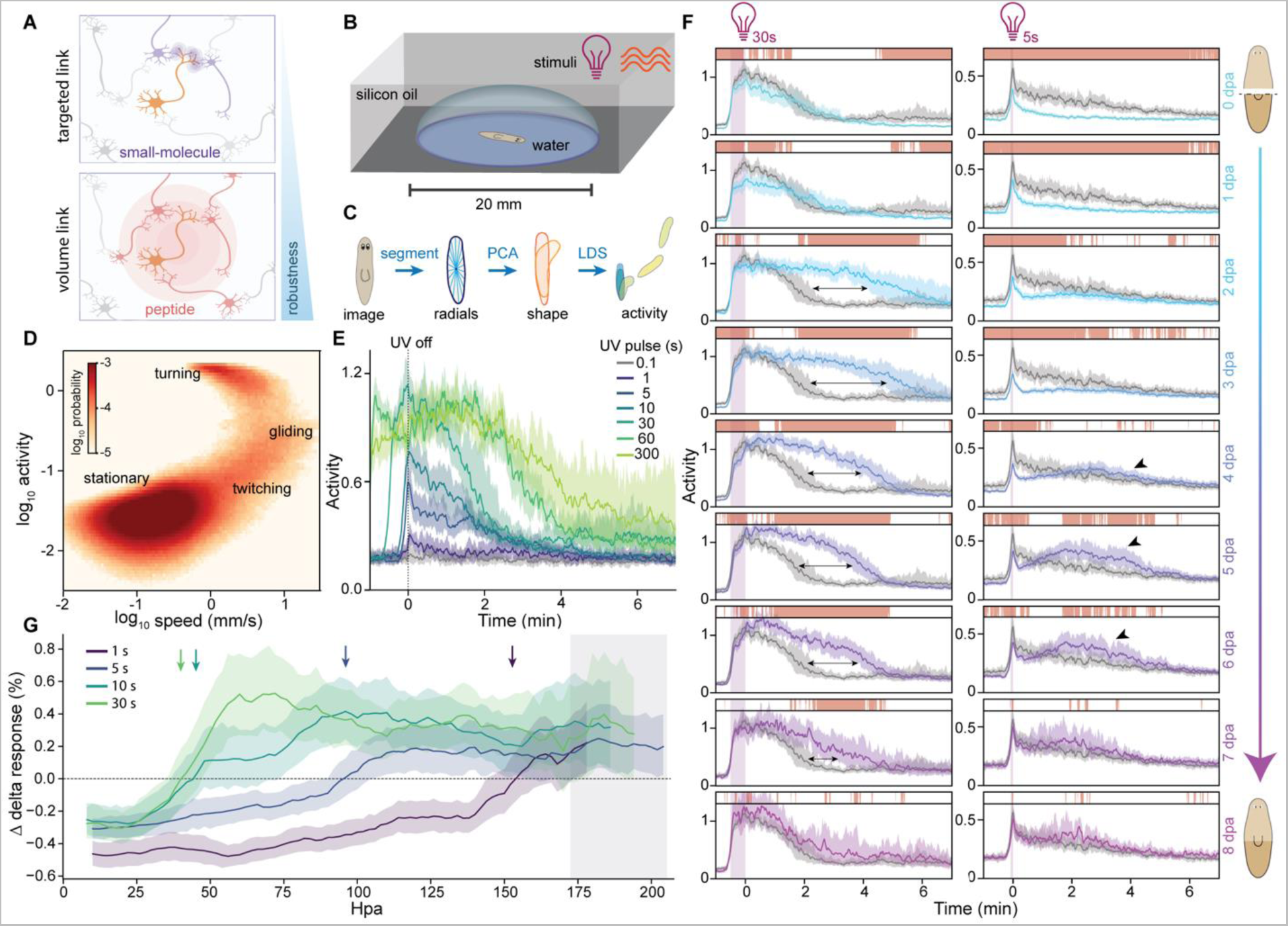
High-content imaging quantifies behavioral changes throughout regeneration. **(A)** Schematic showing the different ranges of peptidergic and small-molecule signals. Neuropeptides create long-range volumetric diffusive cues that can increase robustness without the need to change synaptic network topology. **(B)** Schematic showing the imaging setup with individual planarians recorded in separate aqueous droplets under IR illumination and UV or mechanical stimulations. **(C)** Data processing pipeline includes segmentation, shape quantification using radial measurements, dimensional reduction through principal component analysis (PCA), and a linear dynamics system (LDS) to define the activity measurement. **(D)** Joint distribution of activity and speed calculated from the behavioral data collected on animals under 5 s UV stimulation. **(E)** Activity response of whole animals to UV stimuli ranging from 0.1 to 300 s. Time zero: end of UV stimulation. For all time traces, solid lines: median activity; shaded region: 99% confidence interval (CI). **(F)** Response to UV stimulation throughout the time course of regeneration after decapitation. Gray traces: whole-animal controls; colored traces: regenerating tail fragments. Purple bar: duration of UV stimulation. Arrows: extended response to 30 s UV; arrowheads: ‘resonant peak’ in response to 5 s UV. Bars above the traces indicate time when the response of regenerating animals is significantly different from whole-animal controls as measured by 1,000 nonparametric bootstrap comparisons of the two populations (p < 0.01). dpa: day post amputation. **(G)** Fractional change in UV response during head regeneration relative to the whole-animal control. Average responses are calculated by bootstrap sampling of data from 6 hr before to 6 hr after every time point. Shaded region: 99% CI; arrows: end of reduced activity and beginning of excess activity; grey shaded zone: end of excess activity. hpa: hour post amputation.

We demonstrate neural robustness based on long-range diffusion through both experimental and computational model systems. Experimentally, we study the planarian flatworm *Schmidtea mediterranea*, a basal cephalized animal with the ability to regenerate its entire nervous system from small tissue fragments^18–22^. This regenerative ability is key to the survival and reproduction of planarians, which undergo asexual fission by ripping off tail fragments which then develop into new individuals^23^. The planarian nervous system contains diverse neural cell types and complex structures including a bi-lobed brain, ventral nerve cords, and peripheral projections^24,25^. The number of neurons may fluctuate between ~1,000 to ~100,000 in a single animal during growth and degrowth, requiring dynamic scaling of the entire neural architecture^18^. At the functional level, planarians show complex behavior that integrates information from chemical, light, temperature, and mechanical stimuli^26^. In particular, ultraviolet (UV) light and mechanical cues are detected through sensory cells distributed throughout the body and can stimulate reflex-like responses in decapitated animals^7,27,28^, though it is unclear whether more complex behaviors such as sensory integration are similarly independent of the brain. The cellular and molecular underpinnings of the planarian’s behavioral robustness remain mostly unexplored.

By developing a long-term high-content imaging platform, we observed thousands of planarians during homeostasis and regeneration and quantified their behavior through six orders of magnitude in time. This allowed us to identify previously unknown behavior including signal integration and short-term memory, which revealed a long-lasting latent state in the planarian nervous system. We discovered that maintenance of this excited state is mediated by neuropeptides and demonstrably more robust than inhibition, which is controlled by locally-acting small-molecule neurotransmitters. Using a dual-channel neural signaling model, we show that the different time and length scales of neuropeptide and small-molecule transmission create interfering patterns of neural activity which competitively control neural population dynamics. Though genetic and surgical structural network disruptions perturb both transmission mechanisms, long-range diffusion allows peptide-mediated dynamics to better persist than those driven by small-molecules. This allows peptide function to dominate, generating robust behavioral output. By dynamically balancing contributions of the two signaling mechanisms, this mode of ‘adaptive robustness’ achieves more consistent control after injury than either system would alone.

## Results

### High-content imaging reveals an excitable latent state in the planarian behavior

To uncover complex behavior of planarians such as sensory integration and memory, we imaged freely behaving planarians for extended periods, which has been challenging due to their strong preference for solid edges and photophobic responses^29^. We developed quasi-2D fluidic chambers^30^ to contain animals (**Figure 1B**) and used infrared (IR) for non-perturbative illumination^31,32^. We also incorporated programmable UV (365 nm) and vibrational stimuli to drive ecologically relevant behavior through distinct sensory pathways^26–28,33^. We confirmed that planarians could be continuously imaged on this setup over multiple days without significant changes to their behavior (**Figure S1**).

Continuous imaging with a sub-second resolution allowed us to track animals and quantify the changes in their positions over time (i.e., speed). Additionally, we defined a scalar activity measurement based on the rate of change in the planarian’s shape (**Figure 1C**, see **Methods**). This activity score differentiated gliding, turning, and twitching, whereas speed did not resolve the latter two (**Figure 1D, Supplemental Movie 1**). Activity measurements enabled us to quantify responses with high sensitivity and precision across a broad range of UV doses (**Figure S2**). Beyond known immediate reflexes^7,27^, short UV pulses (<10 s) resulted in exponentially decaying post-stimulus activity, and longer stimulation caused persistent high activity for several minutes before decay (**Figure 1E**). The duration of high activity extended proportional to the dose of stimuli, implicating an ability to both integrate stimulus and differentiate the subsequent behavior through maintenance of activity over several minutes and inhibition of responses at appropriate end points. This led to a broad power-law scaling between pulse duration and total post-stimulus activity, consistent with Steven’s law^34^ (**Figure S2A**).

### Maintenance and inhibition of behavioral state exhibit differential robustness during neural regeneration

This readout of minutes-long memory enabled us to study how planarian’s information processing ability changes during neural injury. We bisected planarians to completely remove the brain and measured the UV response of tail fragments every two hours throughout regeneration (**Figure 1F**). The response to long UV pulses (e.g., 30 s) remained largely intact, though with a reduction in peak and total response after amputation. Strikingly, by 2 days post amputation (dpa), peak activity was not only fully restored, but the response duration, measured as the time needed for activity to return to the baseline, was maintained twice as long after the stimuli compared to whole-animal controls, indicating an inability to properly inhibit behavior output. Excess activity (total activity beyond that of the control) gradually decreased as regeneration progressed and disappeared at ~8 dpa. Response to shorter pulses (e.g., 5 s) followed a similar trend, with initially reduced responses, excess activity appearing as a second ‘resonant’ post-stimulus peak during 4-6 dpa, and full recovery at 8 dpa. Notably, brain regeneration occurs in a similar timeframe, with a primordial brain forming at ~3 dpa and undergoing structural development over the following week^19–21^.

Varying pulse duration between 1-30 s, we found that each stimulus generated reduced responses on the first day after amputation, suggesting impaired ability to maintain post-stimulus activity. Recovery of this ability caused excess activity to appear earlier during regeneration in response to higher UV doses. In contrast, suppression of excess activity did not occur until ~7 dpa, with the timing mostly independent of the UV dose (**Figure 1G, Figure S3**). The fact that maintenance and inhibition of UV-stimulated behavior recover asynchronously in regeneration suggests that they are controlled by separate neural processes which have different capacities to function within a partial nervous system.

To test whether this phenomenon was specific to the UV-sensory circuit, we stimulated planarians using mechanical vibration which is sensed orthogonally to UV^28^. The post-stimulus response to vibration followed a similar dose-response scaling as UV (**Figure S4A,B**). Following amputation, vibration responses showed the same phases of recovery: early in regeneration the response was reduced, then rebounded beyond the whole-animal controls before converging to the baseline (**Figure S4C**), implying that the progression of behavioral recovery is likely governed by changes in post-sensory neurons.

### Peptidergic and small-molecule signals maintain and inhibit behavioral states respectively

We next sought to identify the neural transmission systems controlling the activation, maintenance, and inhibition of post-stimulus behavior. We began by disrupting the core SNARE complex, including *syntaxin*, *synaptobrevin*, and *snap25*, which mediates synaptic vesicle fusion and release^35^. These RNAi experiments all resulted in loss of the UV response (**Figure 2A**), demonstrating that the synaptic network is required for producing behavioral output in planarians. In addition, RNAi of the vesicular glutamate transporter, *vglut,* severely reduced UV response (**Figure 2B**) and caused uncoordinated movement **(Supplemental Movie 2**), suggesting that glutamate is a key synaptic transmitter in planarians.

**Figure 2:**
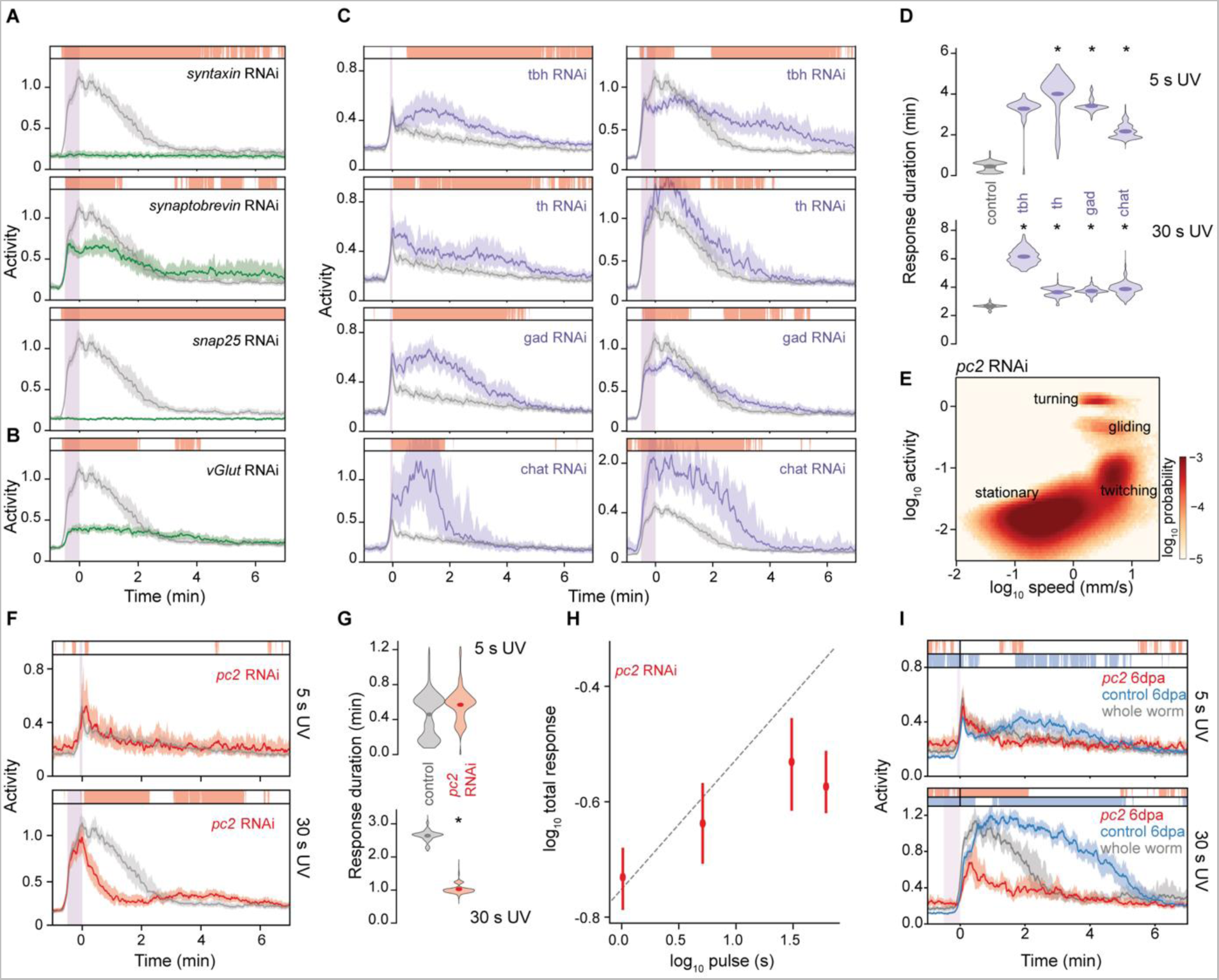
Peptide and small-molecule signals play opposing roles in regulating UV responses. **(A)** Responses to UV stimulation after disrupting components of the SNARE core complex. Purple bar: duration of UV stimulation. **(B)** Responses to UV stimulation after *vglut* RNAi. **(C)** Responses to UV stimulation after disrupting small-molecule neurotransmitter syntheses. **(D)** Duration of post-stimulus activity under RNAi and control conditions. Symbols: mean estimate from 1,000 non-parametric bootstrap samples; histograms: bootstrapped sampling distribution. **(E)** Joint distribution of activity and speed after *pc2* RNAi under continuous UV simulation. **(F)** Left: UV response after *pc2* RNAi. Right: Duration of post-stimulus activity is reduced after *pc2* RNAi. **(G)** Duration of post-stimulus activity under *pc2* RNAi and control conditions. Symbols: mean estimate from 1,000 non-parametric bootstrap samples; histograms: bootstrapped sampling distribution. **(H)** Total response to UV stimulation of *pc2* RNAi animals saturates at high UV dose. Error bars: 99% CI. Line: anticipated power-law relationship (Extended Data Figure 2b). **(I)** UV response in 6 dpa regenerating tails with (red) and without (blue) *pc2* RNAi. Gray: whole-animal response. Statistics. In (A), (B), (C), and (F), gray traces: RNAi controls; colored traces: RNAi of specific genes. Bars above the traces in (A-D), (F), and (I) indicate times when the response is significantly different from controls as measured by 1,000 nonparametric bootstrap comparisons of the two populations (p < 0.01). Asterisks in (B) and (D) indicate p < 0.01, determined by 1,000 non-parametric bootstrap comparisons of the difference in response duration between control and RNAi groups.

We reasoned that regulation of behavioral states on the minute time scale may be governed by neuromodulators shifting the patterns of neural firing. We used RNAi to knock down synthesis enzymes of various small-molecule neurotransmitters/neuromodulators and neuropeptides and found that disruption of octopamine (i.e., tyramine beta-hydroxylase, *tbh*, RNAi), dopamine (tyrosine hydroxylase, *th*), GABA (gabaergic decarboxylase, *gad*), and acetylcholine (choline acetyltransferase, *chat*) syntheses all led to excess post-stimulus activity in response to UV. Similar to the excess activity observed during regeneration, these knockdowns resulted in a second resonant peak in activity after 5 s UV stimulation and a significantly delayed decay of activity after 30 s UV pulses (**Figure 2C,D**). *chat* RNAi also increased peak activity. This suggests that inhibition of post-stimulus activity depends on the cumulative function of multiple small-molecule neurotransmitters.

In contrast, reduction of post-stimulus activity was only observed when knocking down prohormone convertase 2 (*pc2*), which is required for the maturation of many planarian neuropeptides^36^. Planarians have densely packed peptidergic neurons collectively expressing a suite of >60 neuropeptides, most, if not all, of which are also capable of generating small-molecule neurotransmitters^24,36,37^. Long-range peptide transmission could create a densely connected network as almost every planarian neuron expresses some neuropeptides or neuropeptide receptors^24^. While peptides are known to often act synergistically^38,39^, disrupting *pc2* allows for reduction of overall peptide concentrations^36,40^.

Though previous work noted that *pc2* knockdown severely reduces coordinated movement^41^, we found that *pc2* RNAi animals under continuous UV stimulation could activate the full range of behavior seen in control animals (**Figure 2E, Supplemental Movie 3**). Consistently, their response to UV stimuli was activated to levels matching that of controls and showed little differences with weak stimuli. However, unlike controls that maintained high activity after long stimulation (30 s), responses in *pc2* knockdown animals decayed immediately (**Figure 2F,G**). This prevented the animals from differentiating their responses to long stimuli and caused saturation in the dose-response curve (**Figure 2H**). These observations suggest that, while *pc2* RNAi animals can detect and respond to UV, they fail to integrate signals and maintain the latent memory state needed for extended post-stimulus activity. Concordantly, when we amputated *pc2* knockdown animals, they failed to show either an extended response or resonant peak after 30 s and 5 s UV pulses, respectively, during regeneration (**Figure 2I**). Altogether, these results suggest that long-range peptide signaling underlies the rapid return of response maintenance after injury, whereas small-molecule signals mediate the more fragile inhibitory functions.

### Neuropeptide signaling maintains short-term memory

We hypothesized that other forms of memory at this timescale may also be mediated by peptides and similarly robust to injury. To test this, we exposed planarians to pairs of 5 s UV pulses separated by a time delay. With delays on the order of minutes, the response to the second pulse is significantly stronger than that of the first (**Figure 3A**), demonstrating sensitization, a form of short-term memory. This could be enhanced by stronger first pulses (**Figure 3B**). To measure how the memory of the first pulse changes over time^42^, we varied the delay between pulses and found a non-monotonic decay of the sensitizing effect with a secondary peak at ~ 3 min delay (**Figure 3C**). Sensitization is also seen when pairing mechanical vibration and UV pulses, suggesting that this memory is embedded in post-sensory processes (**Figure 3D**).

**Figure 3:**
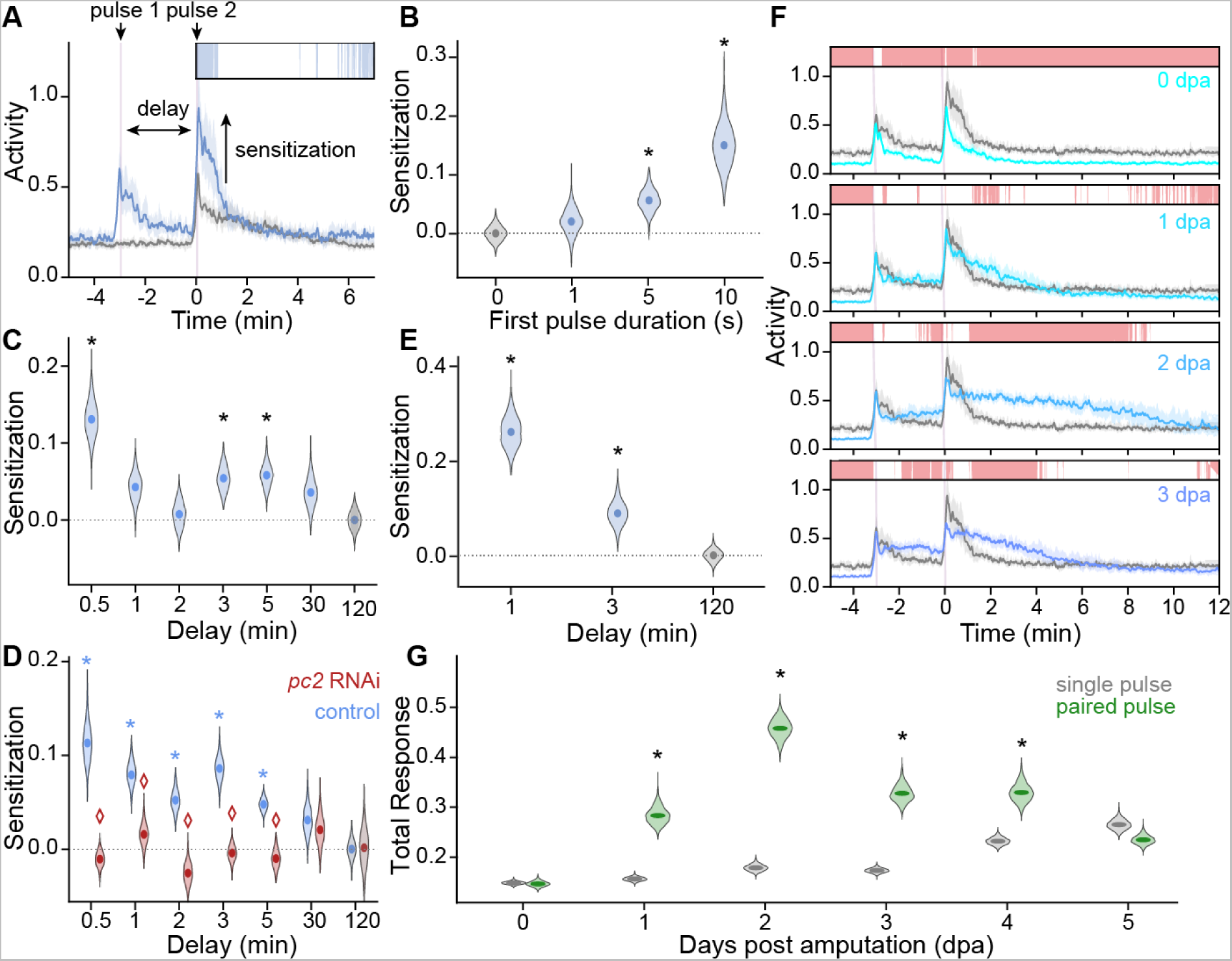
Neuropeptides are required for establishing short-term memory. **(A)** Planarians show sensitization from prior UV exposure. Gray: response to single 5 s UV pulse; blue: response to paired 5 s pulses separated by 3 min delay. For all time traces, lines: median activity; shaded region: 99% CI. **(B)** Sensitization, defined as the difference between total post-stimulus response to the second pulse and that of a single UV pulse after a 3 min delay, increases with the duration of the first UV pulse. **(C)** Sensitization from two 5 s UV pulses vs. delay time. **(D)** Sensitization is lost in *pc2* RNAi (red) animals. **(E)** Mechanical vibration (5 s) sensitizes response to 5 s UV pulse. **(F)** Response of paired 5 s UV pulses with 3 min delay through regeneration. Gray traces: whole-animal controls; colored traces: regenerating tail fragments. Purple bar: duration of UV stimulation. **(G)** Total response in tail fragments to a single 5 s UV pulse (gray) and a 5 s pulse sensitized by another 3 min prior (green) shows that the sensitization amplifies at early time points of regeneration. Statistics. Bars above the traces in (A) and (F): timepoints where activity in paired pulse is significantly different from that of single pulse as measured by 1,000 nonparametric bootstrap comparisons (p < 0.01). For all violin plots, symbols: mean estimate of sensitization using 1,000 non-parametric bootstrap samples of both the paired and single pulse conditions; histograms: bootstrapped sampling distribution. Asterisks: sensitization significantly greater than zero (p < 0.01); diamond: significant difference in sensitization between control and *pc2* RNAi (p < 0.01).

While *pc2* RNAi did not affect response to single 5 s UV pulses (**Figure 2E**), it eliminated sensitization, indicating that neuropeptides are required for maintaining short-term memory (**Figure 3E**). Sensitization was initially lost in amputated planarians, but rapidly increased beyond that of whole animal controls at 1 dpa, demonstrating rapid recovery of sensitization memory but a lack of inhibition (**Figure 3F,G**). This observation parallels the trend in the single-pulse responses during regeneration, suggesting that the same peptide-dependent excitable latent state may encode both signal integration and short-term memory.

### Peptide mediated functions are more robust to general brain perturbations

If long-range neuropeptide transmission is less reliant on the intact network structure than small-molecule signaling, then lost inhibition and excess behavioral activity should be a generic signature of brain injuries. To test this prediction, we performed several surgical cuts causing partial brain damage, including severing anterior commissures (i.e., ‘corpus callosum cut’), amputating anterior to the eyespots, and biopsying posterior to the left eyespot. Even though these injuries affected different neural structures, they all led to similarly extended UV responses within the first day after injury (**Figure 4A**).

**Figure 4:**
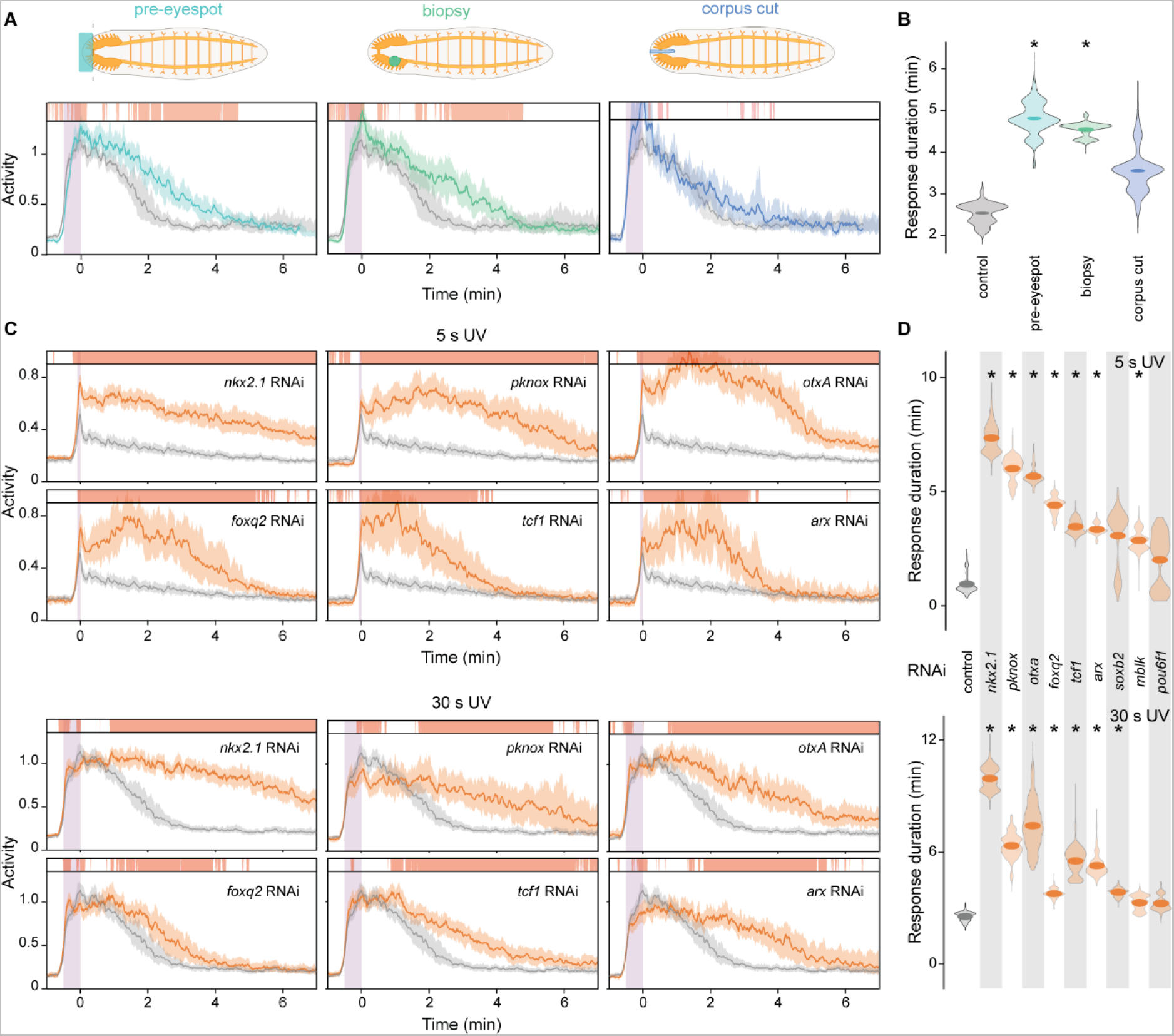
Excess activity is a signature of perturbed neural structure. **(A)** Response to 30 s UV shows extended activity after various minor injuries. Colored traces: injured animals at 1 dpa; gray traces: whole-animal controls. Purple bar: duration of UV stimulation. **(B)** Violin plot showing the duration of post-stimulus activity. Symbols: mean estimate from 1,000 non-parametric bootstrap samples; histograms: bootstrapped sampling distribution. Asterisks: p < 0.01 comparing injured and whole-animal responses. **(C)** Response to 5 s (top) and 30 s (bottom) UV after knockdown of neural TFs. Orange traces: RNAi; gray traces: control. **(D)** Response duration after RNAi. Asterisks: p < 0.01 comparing RNAi and control conditions. Statistics. Bars above the traces in (a) and (c) indicate the time when the response is significantly different from whole-animal controls as measured by 1,000 nonparametric bootstrap comparisons of the two populations (p < 0.01). Solid lines: median activity; shaded region: 99% CI; p-values in (B) and (D) are determined by 1,000 non-parametric bootstrap comparisons of the difference in response duration between the two groups.

To rule out the possibility that excess activity is driven by wound response, we also perturbed neural structures genetically by performing RNAi to disrupt a set of 9 transcription factors (TFs) known to play important roles in the development of various neuronal populations and other processes during brain regeneration such as patterning and size regulation^20–22,24,43^. Despite the distinct functions of these TFs, almost all knockdowns led to similar excess activity in response to UV through extended durations and higher activity peaks without physical injury (**Figure 4B,C, Figure S5**). The strikingly consistent effect across surgical and genetic perturbations implies that the differential robustness of maintenance and inhibition of the latent state is likely not caused by asynchronous regeneration of controlling neural populations. Instead the two processes may be encoded through distinct patterns in population-scale neural dynamics, with peptide-mediated dynamics more robust to structural changes independent of specific neural circuits.

### Long-range volumetric transmission explains the robustness of peptide signaling

We then asked whether differential signaling ranges are sufficient to explain the observed difference in the robustness of peptide and small-molecule mediated processes. To do so we developed a neural network model in which neurons interact through both volumetric and synaptic signals and constrained the two systems to regulate behavior in the same manner seen in planarians. We then tested whether these mechanistic underpinnings were sufficient to stabilize peptide communication and generate excess behavioral output upon neural injury.

Specifically, we modeled population-scale neural dynamics using a custom recurrent neural network (RNN) which simulates neuronal firing coupled through sparse synaptic links and produces behavioral output through a readout matrix applied to the firing rates^44^. This model has been previously shown to reproduce key aspects of animal neural dynamics when trained on similar tasks^44–46^. To account for long-range volumetric signals, we augmented the model by allowing neuron firing to trigger release of excitatory and inhibitory neuropeptides. These peptides diffuse and persist extracellularly and modulate neuron firing potentials based on local concentrations (**Figure 5A, Figure S6A**, see **Methods** for model details).

**Figure 5:**
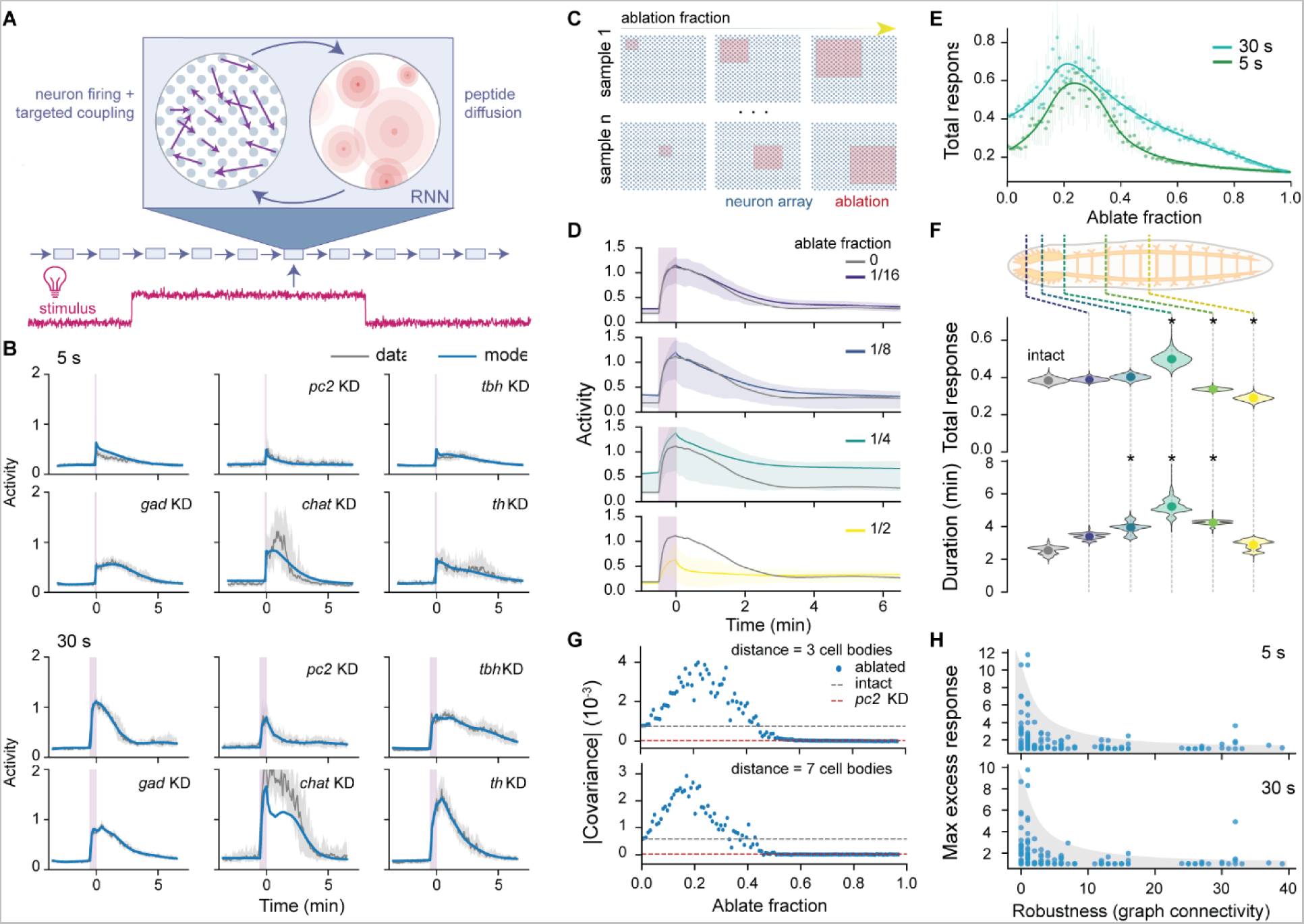
A RNN model recapitulates functional robustness after neural injury. **(A)** Schematic of the dual-channel RNN model showing the feedback between peptidergic and small-molecule signaling. **(B)** Output of model trained to emulate planarian responses under various stimulation and knockdown (KD) conditions. Gray: experimentally measured median activity; shaded region: 99% CI; blue: model response. **(C)** Schematic of neural ablation within the model. Red regions are spatially contiguous populations of neurons with firing rates fixed at zero. For each ablation fraction, 200 regions are randomly sampled. **(D)** Model response to 30 s UV stimulation at different ablation fractions. Gray: response of intact model; colored: mean activity over 200 sampled ablations; shaded region: interquartile value of activities across ablation samples. **(E)** Total response after ablations shows excess activity at moderate ablation fractions. Symbols: average total response across ablations; error bars: 99% CI from 1,000 non-parametric bootstrap samples; solid lines: polynomial interpolation. **(F)** Planarian response to 30 s UV shows similar trends with increasing ablation as in the model. Landmarks for amputation (top): anterior-end of eyespot, posterior end of eyespot, one eyespot length posterior to eyespot, midway between eyespot and pharynx, mid-pharynx. Data for all injured conditions are collected at 1 dpa. Symbols: mean estimate from 1,000 non-parametric bootstrap samples; histograms: bootstrapped sampling distribution. Asterisks: p < 0.01 comparing amputated and intact worms. **(G)** Magnitude of spatial covariance of firing rates, which provides a quantification of the relative contributions of the two communication mechanisms. Expectedly, the spatial covariance was lost with *pc2* knockdown, but amplified when reducing small-molecule signals (Extended Data Figure 6b). Distances at which the covariance is measured correspond to the diffusion length scales of the two peptides in the trained RNN model. **(H)** Ablation-induced excess activity diminishes with increasing connectivity network robustness quantified by the graph connectivity. Each dot represents an independently trained RNN with different connectivity networks. The maximum excess response is defined as the ratio of maximum average total response at any ablation level and the total response in the intact model.

Because the model is implemented in a deep-learning framework, we were able to constrain its dynamics to produce experimentally observed behavior. Simply training on wild type response is insufficient to ensure that long-range peptide and targeted small-molecule signals in the model maintain and inhibit behavioral activity, respectively. Therefore, we also trained on *pc2* RNAi data while blocking peptide transmission in the model, allowing it to learn the effective contribution of the volumetric signal. Similarly, we trained on data of RNAi blocking small-molecule transmission to constrain their functional roles. Because of the relatively fast and targeted scales of small-molecule transmission, we model their contribution as instantaneous action through synaptic links and assign subsets of neurons as octopaminergic, dopaminergic, GABAergic, and cholinergic. When training on the RNAi data, transmission of the corresponding small molecule was blocked while allowing neural firing and peptide release to sustain. This protocol was sufficient to train models that each fit all the experiments simultaneously (**Figure 5B**).

We then tested whether these constraints were sufficient to explain the differential robustness of the two communication mechanisms. We simulated the effects of neural injury by ablating regions of neurons at various locations and sizes (**Figure 5C-E**). Matching experiments, large ablations reduced the response, corresponding with early regeneration time points after decapitation when the network is highly disrupted. Moderate ablations led to an extension of the response like those observed in partially regenerated animals (**Figure 1E-G**). For a more direct experimental validation, we amputated planarians to remove increasingly larger anterior structures and measured response to 30 s UV pulses at 1 dpa. We observed the same trend of progressive increases of duration and total responses before loss of response with largest amputations (**Figure 5F**). This consistency between experiments and model predictions demonstrates that the robustness of peptide signaling and the early return of peptide-mediated maintenance of behavioral state after injury can be explained solely by the difference in transmission mechanisms, regardless of the circuit details.

### Adaptive robustness through incoherent signal competition

This dual-channel model also allowed us to study why maintenance and inhibition of behavior segregates by mechanism of neural transmission. For this, we measured how the two mechanisms interact to drive neural population dynamics. Due to long decay time and long-range diffusion, peptide signaling generates slowly varying, spatially correlated patterns of neural activity. In contrast, targeted connections and local neural modulations produce rapidly changing, spatially uncorrelated neural firing patterns. Because both mechanisms are broadly co-expressed in planarian neurons^24^, these different patterns of neural activity propagate in a shared medium. Mechanistic differences of the two systems create incoherent correlation structures which effectively compete for control over neural population dynamics. By aligning motor output to components of neural dynamics driven by peptide signaling (e.g., spatially continuous patterns), peptide function should stabilize and maintain behavior activity, while high-frequency targeted signals disrupt these patterns, drive the neural activity to new states, and effectively inhibit behavior activity (**Figure S6B**).

Upon ablation, propagation of small-molecule mediated signals were more severely disrupted, reducing their competitive influence on firing patterns and increasing the relative contribution of neuropeptides. This effect could be quantified by the peptide-mediated spatial correlation in firing dynamics, which became more pronounced after moderate ablations, explaining the excess behavior seen in this regime (**Figure 5G**). This creates a system of behavior regulation through neural pattern competition that is largely independent of specific neural circuitry, providing a new paradigm of adaptive robustness during massive structural changes.

To further validate that differential robustness between signaling mechanisms drives the adaptive robustness phenomenon, we varied the topological robustness of synaptic connectivity by training over 150 independent RNN models with different connectivity network structures (**Figure S6C**). More robust synaptic networks maintained their contribution to neural dynamics and prevented ablation-induced excess activity even though the peptide transmission was kept identical (**Figure 5H**). This provides direct evidence that the excess activity induced by neural injury is caused by the different capacity of the peptidergic and small-molecule signaling to maintain their functions in disrupted neural networks.

## Discussion

In this study, we used the planarian’s ability to regrow an entire head de novo to study the neural processes underlying its robust behaviors during brain regeneration. Extensive behavioral data and computational analysis using a dual-channel signaling network led us to a conceptual model of adaptive neural robustness (**Figure 6**). During structural injury, both channels are perturbed in a correlated manner, making their competitive output, i.e., difference in strength, more robust than the function of either alone. As the animal regenerates after injury, peptide-based functions recover quickly due to the capacity of long-range diffusion to cross disrupted regions. In contrast, targeted small-molecule functions require more complete connectome and thus take much longer to return. By aligning behavioral output with the more robust peptide-dominant patterns of neural activity, the planarian ensures reliable motor sensory responses throughout massive neural remodeling.

**Figure 6:**
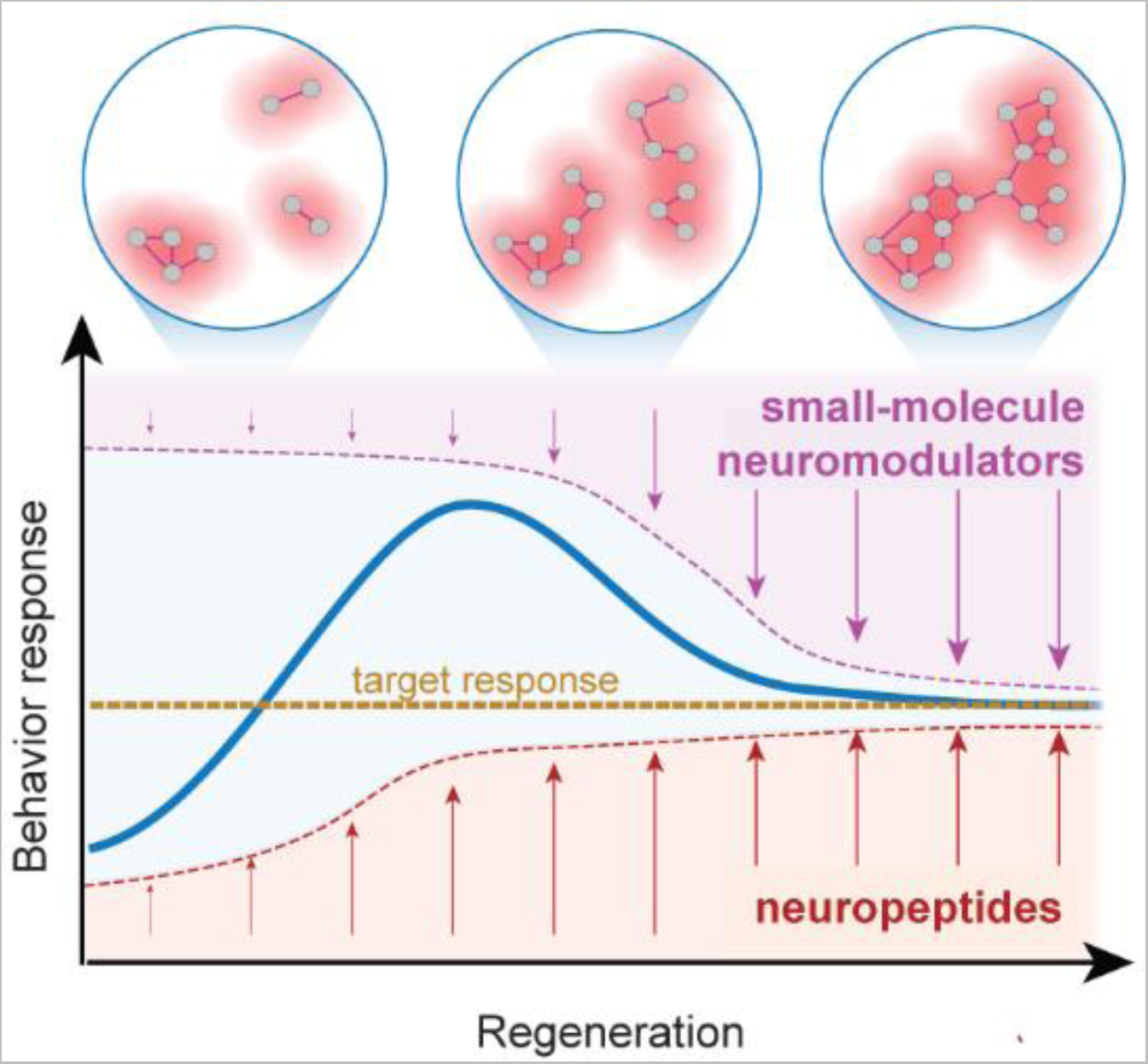
Adaptive robustness through multi-channel signaling. Long range peptide signals (red) and targeted small-molecule signaling (purple) form a dual-channel network and competitively regulate behavioral output (bottom). Under highly disrupted post-amputation conditions (top left), the functions of both systems are compromised. As regeneration proceeds, peptides can rapidly establish communication via long-range volumetric diffusion to drive behavioral output, while the fragile targeted network remains fragmented dysfunctional (top middle). The transient dominance of peptidergic signals at this stage leads to excess activity in response to stimuli, which is then gradually refined as the connectome and targeted small-molecule signaling re-establishes. When regeneration is complete, both transmission mechanisms are restored and the behavior is constrained to the proper response (top right).

In contrast to neurogenesis during embryonic development that occurs in protected environments such as an egg, neural regeneration must proceed while responding to threats from the surrounding world. For example, protection from UV irradiation is critical as it induces DNA damage, which can become particularly harmful during regeneration due to elevated cell proliferation^47^. Allocating functions activating strong UV responses to a relatively robust communication mechanism allows for rapid re-establishment of key survival mechanisms like the escape response while permitting other features to be re-established later. Short-term memory also provides a means for animals to evaluate changes in stimuli across space and time. This can enable phototaxis at length scales much larger than their body size^48,49^, which is essential for planarians to locate areas of lower exposure to minimize damage induced by UV. Our results demonstrate that short-term memory is also mediated by neuropeptides allowing for recovery very early during regeneration. Together, our findings imply that mechanisms enabling rapid behavioral recovery after injury can provide an advantage under selective pressures in regenerating animals, especially in organisms that reproduce through repeated fission and regeneration.

The long-range diffusion and slow time scales of neuropeptide signaling offer a molecular basis for long-lasting latent neural states^50,51^, which need to be robust after injury in order to properly process information and maintain behavioral output. Indeed, the usage of peptide signals to stimulate behaviors and promote arousal states has been observed in diverse animals including nematodes, flies, and even mammals^15,52–54^. In some cases, it has been shown that the peptide-dependent circuits function opposingly to synaptic connectome such that the interactions between the two drive the switching between behavioral states^15,55^. The parallel across different organisms implies that the division of labor between the two transmission mechanisms may be a common feature of many neural circuits. Examining whether these circuits possess similarly high robustness is an important avenue for future research.

## STAR Methods

### RESOURCE AVAILABILITY

#### Lead Contact

Further information and requests for resources and reagents should be directed to and will be fulfilled by the lead contact, Bo Wang.

#### Materials availability

This study did not generate new unique reagents.

#### Data and code availability

All activity data used for analysis and model training are available at tinyurl.com/robustBehavior. Original videos and radial segmentation measurements are available from the authors upon request, due to the lack of public repository to host such large volumes of data.

Software for video segmentation is available at github.com/samuelbray32/planameterization. Repository for data analysis and visualization is available at github.com/samuelbray32/PARK. Repository for dual-channel RNN model construction and training is available at github.com/samuelbray32/dualChannelRNN.

### EXPERIMENTAL MODEL AND SUBJECT DETAILS

#### Animals

Asexual *S. mediterranea* were maintained in the dark at 20 °C in water supplemented with 0.5 g/L Instant Ocean Sea Salts (Carolina Biological Supply, Cat#671442) and 0.1 g/L sodium biocarbonate. Planarians of ~4 mm in length were used for whole-animal behavior experiments and were fed once or twice a week. For amputation experiments, we selected animals of ~8 mm in length such that the regenerated fragments were approximately the same size as those used in whole-animal experiments.

#### RNAi

Gene knockdowns were carried out by feeding double-stranded RNA (dsRNA) to induce RNAi mediated gene silencing. For all experiments, we fed dsRNA to animals 5-7 times every 4-5 d, except *pc2* for which we fed 3 times. Animals were starved for 4 d prior to amputation, after which tails were allowed to regenerate and imaged at 10-20 dpa, except for *pc2* RNAi animals, which were imaged 5 d after the last feeding without amputation. Knockdown of *synaptobrevin* and *syntaxin* caused lysis after ~20 dpa, but no gross morphological phenotypes were observed during the time window of imaging.

The dsRNA was synthesized using the established protocol^36^ and fed to animals by mixing in a liver homogenate at a concentration of approximately 100 ng/µl. All clones for dsRNA synthesis were generated using oligonucleotide primers (**Supplemental Table 1**) and cloned into vector pJC53.2 (Addgene plasmid ID: 26536)^36^. For the RNAi control condition, we fed dsRNA matching the ccdB and camR-containing insert of pJC53.2 under the same schedule of *pc2* RNAi. Results from these experiments showed no significant difference from animals without RNAi feedings. To maximize statistical power, we therefore combined data from these two conditions as the control to compare against all other knockdown conditions.

### METHOD DETAILS

#### Imaging chamber

To create the imaging chambers, 150 mm petri dishes were plasma treated (Harrick Plasma, PDC-001) for 2 min at high power to create a hydrophilic surface. Ten circular templates with 20 mm diameter were traced in each dish with a permanent marker to pattern a hydrophobic mask. Each region was first loaded with a single planarian in 800 µl of Instant Ocean water to ensure complete wetting and then reduced to ~350 µl. This flattened the top surface of the droplet and reduced reflections during imaging. After loading, the dish was gently filled with ~50 mL of silicone oil (Fisher Scientific S159-500) until the top of all droplets were covered. The oil phase both reduced evaporation and increased surface tension at the droplet boundary to prevent planarians from escaping. Planarians remained viable and active for at least ~7 d at 20 °C under these conditions. All animals were starved at least 4 d prior to imaging and all imaging sessions lasted no longer than 7 d. For imaging regeneration time courses, animals were loaded at 8 hpa and rinsed before loading to reduce the accumulation of material ejected from the wound site within the droplet. To cover the complete span of regeneration, a second cohort of animals was imaged starting 3 dpa for ~7 days. Activity responses at matching regeneration time were merged from both datasets and used for all analyses.

#### Imaging setup

Animals were illuminated with an IR light source (850 nm) from the side, and images were recorded using a Rasberry Pi NoIR camera, except for a higher resolution camera (Daheng Imaging, MER-1220-32U3M) used to generate the movies shown in **Supplemental Movie 1-2**. UV light (365 nm) stimuli was delivered by a custom-built ring of 36 LEDs (Waveform Lighting, 7021.65) mounted above the camera to illuminate the entire dish uniformly and controlled by an Arduino Uno (A000066) to adjust intensity and pulse duration. To eliminate glare from the stimulus light source, an 800 nm long-pass filter (ThorLabs, FELH0800) was mounted within the camera tube. Tactile stimulation was delivered by vibrating the stage with a small motor (Vibronics Inc, VJQ24-35F580C), which was also controlled by the Arduino. For all stimulation experiments, repetitions of the stimulation protocols were separated by 2 hr of unstimulated time to prevent influence between trials.

#### Dual-channel RNN model

We modified a well-established RNN model^44^ by adding long-range volumetric signals. All neural models were implemented, trained, and run using Keras architecture with Tensorflow 1.14 backend in a Python environment with CUDA GPU acceleration^56^. For a graphical diagram of model architecture, see **Figure S6A**.

Models trained within this work contain 2500 neuron nodes arranged in a 50 × 50 square grid (*n_neuron_*) and 2 neuropeptides (*n_pep_*). These values were chosen to maximize flexibility in model training while maintaining loss gradients that could be stored in working memory. The RNN cell has 5 state tensors passed between timepoints: a *n_neuron_* vector of neuron states (*X_syn_*), an array of each neuropeptide concentration at each neuron location (*X_pep_*), a binary variable indicating whether neurons are capable of responding to neuropeptides in the sample condition (*𝑔_pep_*), a vector describing the modulatory potency of each neuron in the sample condition (*𝑔_syn_*), and a vector describing the ablation status of each neuron (*A*).

The model has one fixed parameter, a *n_neuron_* × *n_neuron_* connectivity matrix (*C*), which is a sparse, binary matrix. This matrix is defined using the Watts-Strogatz method^57^. By varying the average node-degree (*k*) and rewiring probability (*β*), this method can create networks with topologies ranging from a completely regular ring structure (*β* = 0), to small-world networks (0 < *β* < 1), to a random network (*β* = 1). As previous studies have demonstrated ‘rich club’ small-world networks in a variety of neural systems^58^, we defaulted to using a network with *k* = 8, *β* = 0.001. For the models shown in Figure 5h, triplicate models with different initializations and *C* were trained with node degrees logarithmically spaced from 2 to 64, and *β* logarithmically spaced from 0 to 1 to average the effects of the sparse weighted connections. To compare the robustness of the synaptic connectivity topology, we calculated the graph connectivity of the network using the python package NetworkX^59^. This measure describes the smallest number of nodes that must be removed to separate the graph and corresponds with our simulated perturbation of node ablation.

The RNN cell has 7 trainable model parameters: a *n_simulus_* × *n_neuron_* input vector transforming stimulus condition into influence on the neuron states, a *n_neuron_* × *n_neuron_* weight matrix (*W_syn_*), a decay rate of neuron potential (*δ_neuron_*), the production rate of each neuropeptide upon neuron firing (*μ_pep_*), the decay rate of each extracellular neuropeptide (*δ_pep_*), the diffusivity of each neuropeptide (*D*), the scaling coefficient of peptide action on local neuron potential (*W_pep_*), and a *n_neuron_* vector *W_in_* which defines the transformation of the current stimulus state (*U_t_*) into change in neuron potential.

At each time step, firing rates are sampled from the neuron state according to:

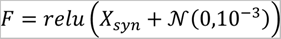

The non-linear rectifier maps the state of the cell into a positive firing rate^44^. Extracellular neuropeptides are updated according to:

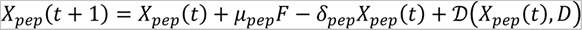

where 𝒟 applies a diffusivity kernel:

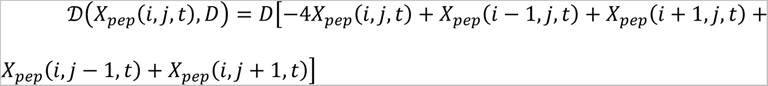

to the peptide concentrations according to their arrangement on the square array of neurons. For calculating spatial diffusion, the array is treated with periodic boundary conditions to remove the effect of system size. Neural state updates are given by:

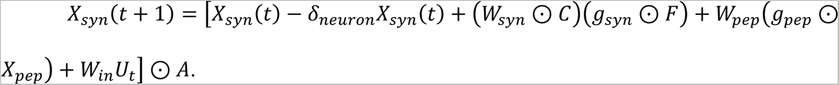

Multiplication of modulatory weights and connectivity enforces a specific, sparse network of connections throughout training. Multiplication of *𝑔_syn_* and the firing rate masks output from genetic knockdown of small-molecules. Multiplication of *𝑔_pep_* and *X_pep_* masks contribution of neuropeptide signaling. Multiplication of the entire state with *A* completely masks all contributions of ablated neurons in a sample.

Model inputs are the stimulus vector across the simulation time range *U*, *𝑔_syn_*, *𝑔_pep_*, and *A*, which are concatenated in the model with *X*_0_, a trainable variable providing the neuron and peptide states at time zero within the RNN. This concatenated tensor is fed as initial conditions to the RNN. The RNN then simulates 1,200 timesteps (10 min) of zero-stimulus response. This allowed the neuron states in a sample to evolve to form a stable state, which is given the knockdown and ablation input. The final equilibrated state is then passed with *U* to the RNN, which simulates the dynamics and returns the complete neural state at each timepoint. This is rectified into the firing rate, and convolved with a single time step kernel, which effectively applies a trainable output matrix *W_out_* to the firing rates at each time step to generate the simulated activity (*Z*). *Z* is returned as the output of the complete machine learning model.

For initialization, elements in large weight matrices were independently sampled according to: *W_syn_*(*i,j*) ~ 𝒩 (2 × 10^−5^, 2 × 10^−4^), neural states of *X*_0_(*i,j*)~𝒩(0,10^−2^), peptide states of *X*_0_(*i,j*)~*Exp*(10^−2^), *W_in_*(*i,j*)~𝒩 (0,5), and *W_out_*(*i,j*)~𝒩(0,0.05). Other parameters were set to: *μ_pep_*= [1,1], *δ_pep_* = [0.2,0.2], *D* = [3,10], *W_pep_* = [0.9, −0.6], *δ_neuron_* = 1.25. These values were chosen to produce non-diverging neural dynamics from which to begin training.

Models were trained on the median activity. To prevent overfitting to the timing of the UV pulse during the simulation, the training data contained 20 samples for each stimulation and knockdown condition with start times randomly selected between 1-4 min before the start of stimulation and a sample duration of 26 min (except for Era 1, in which duration was set to 13 min to reduce size of gradient operations and speed the initial training to obtain a rough shape of the response). The training was performed without ablation (*A* = 1). In samples of *pc2* RNAi data, *𝑔_pep_* = 0. To account for small-molecule neurotransmitter knockdowns, neurons were randomly assigned as GABAergic (12.5%), octopaminergic (12.5%), dopaminergic (12.5%), or cholinergic (50%), or generic (12.5%). The relatively high fraction of cholinergic neurons was chosen to match their abundance in the planarian nervous system^24^, but the model result was only weakly dependent on the neuronal type fractions. We did not specify spatial patterns for these neuronal types to keep the model general and mostly species agnostic. In samples of *gad*, *tbh*, *th*, or *chat* RNAi data, *𝑔_syn_* was set to zero for the corresponding neurons to prevent their small-molecule modulatory activities.

The loss (ℒ) was defined as the mean square error between the median activity and simulated activity Z, and was optimized using ADAM gradient descent^60^. Models were trained in 4 eras:

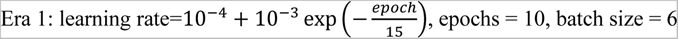

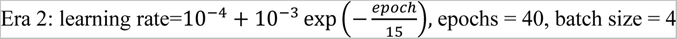

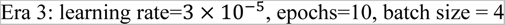

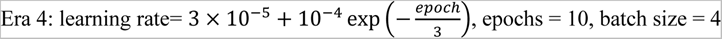

In Era 4, training was stopped early if the change in average loss was less than 5 × 10^−4^ for 3 consecutive epochs.

Each model was trained using responses to 5 s and 30 s pulses for control animals, as well as *pc2*, *gad*, *tbh*, *th*, and *chat* RNAi animals. To further constrain the functional fitting within the model, we performed double knockdowns of *gad:pc2*, *tbh:pc2*, *th*:*pc2*, and *chat:pc2* and included them within the training set. Additionally, to stabilize the long-term behavior of the simulated model, we stimulated animals with UV for 30 min at reduced intensities and included this data within the training set.

After training, the rate parameters in the model shown in detail in **Figure 5** were: D: [2.9952657 10.002745]; *μ_pep_* : [1.006925 0.99327475]; *δ_pep_* : [0.2064426 0.19140989]; *W_pep_* : [0.90302687, −0.59036015]; *δ_neuron_* : [1.2581608]. These parameters gave the range of peptide diffusion to be 3-10 cell bodies and their half-life to be ~ 5 min, which are biologically realistic based on previous measurements^38^.

To simulate responses after ablation, the ablation vector (*A*) was generated to mask out neurons corresponding to a spatially contiguous rectangular region of minimum aspect ratio with the desired fraction of neurons. The location of the ablated region within the array was randomly chosen for each sample. For each ablation fraction, responses were generated from 200 different ablated regions and averaged to remove dependencies on locations of the ablation.

The spatial covariance was calculated by averaging over all pairs of neurons at a given distance and across all timepoints on simulations lasting 11 min with a single 3 s pulse. This was sufficient to drive neural activity without dominating the signal with spatial correlation embedded within the input matrix *W_in_*. Ablated neurons were excluded from the calculation. For each ablation fraction, we simulated results from 200 ablated regions and averaged covariances at each distance.

### QUANTIFICATION AND STATISTICAL ANALYSIS

#### Behavioral activity quantification

Planarians were segmented using binary thresholding. For each frame, the center of mass (COM) of the animal was determined and the perimeter pixel locations were identified and interpolated to give 100 evenly spaced anchor points. Radial distances were calculated as the distance from the COM to each of these points. The radial measurement vector was L1 normalized and aligned such that the maximum radial distance is at position 0. We then reduced the dimensionality of the radial measurements using Principal Component Analysis (PCA) and used the first 10 PCs (97% explained variance) for further analysis^61^. PCs calculated using data from intact animals responding to 5 s UV stimulation were used to analyze all datasets. These PCs correspond to ‘eigen-shapes’ of the worm during movement, e.g., elongation (PC1), turning (PC2), and scrunching (PC3)^61^. For PCs that are symmetric across the worm (e.g., turning left and right gives positive and negative values on PC2 respectively^61^), we took the absolute value of the component to prevent unnecessary separation of similar behaviors.

Because stationary and moving animals occupied overlapping shape space, incorporating temporal information into the feature vector significantly improved our ability to quantify behavioral activity. To do so, we convolved the shape measurement with Haar wavelet filters^62^ at scales corresponding to 2.5, 7.5, and 15 s to obtain the rate of shape change at each time point. This also served to filter out noise from the segmentation steps. Finally, we fit a one-dimensional stochastic linear dynamic system (LDS) model to the wavelet filtered data^63^ using the state space models (SSM) Python package^64^. This model is a continuous analogue of the discrete Hidden Markov Model and infers a single latent scalar variable (activity) at each timepoint which maximizes the likelihood of the observations (wavelet filtered data).

#### Statistical analysis

Planarians displayed an asymmetric multimodal distribution of activities at any given time point. Therefore, we used non-parametric bootstrap statistics to test for significant differences between population averages. When comparing time trace data, each bootstrap sample was performed by choosing *n* responses with replacement from the collected data, where *n* is the number of responses within the given dataset (**Supplemental Table 2**). This created new samples matching the size of the experimental dataset^65^. The median value from these samples at each timepoint was recorded. This procedure was repeated >1000 times to form a sampling distribution of the median population activity at each timepoint from 5 min before to 30 min after the stimulus. All time traces plotted are the mean estimate of median activity with 99% confidence intervals (CIs).

To test for significant differences between two conditions (e.g., RNAi), paired sets of samples were taken from each dataset and the difference in median activity between each sample was recorded. This procedure was repeated to form a sampling distribution of the difference in median activities at each timepoint. Significance at a time point was declared if zero falls outside the 99% CI.

#### Response measurements

To compare responses between conditions, we used three summary statistics. First, total response was quantified as the average median population activity during 10 min post-stimulation. This interval was sufficiently long to capture the dynamics of extended responses while maintaining sensitivity to short pulses. Post-stimulus response duration was defined as the time at which the median activity first fell below a threshold value, 0.3, after stimulation. This threshold was chosen to lie above the confidence interval of unstimulated population activity. Peak response was quantified by the maximum median activity after the end of stimulation. To compare these values between conditions, we used non-parametric bootstrap estimation. We calculated the measurement on 1,000 paired sample sets from control and experimental conditions and recorded the difference. Significance was declared if zero difference fell outside the 99% confidence interval for the sampling distribution of differences.

## Acknowledgements

We thank U Alon, P Nuyujukian, S Granick, and L Luo for critical discussions and H Li for technical assistance. LSW and SRB acknowledge the support from a NIH cellular, Biochemical, and Molecular Sciences (CMB) training grant (T32GM007276). LSW and CC are supported by NSF GFRP fellowships. CC is also a Stanford Graduate Fellow. MEL acknowledges the SSRP-Amgen Scholars Program which provides undergraduate students opportunities to perform research at Stanford University. BW is a Beckman Young Investigator. This work is supported by an NIH grant (1R35GM138061) and the Neuro-omics project of Wu Tsai Big Ideas in Neuroscience program.

## Author contributions

Conceptualization: SRB, LSW, and BW; Methodology: SRB, LSW, CC, and MEL; Investigation: SRB and LSW; Formal analysis: SRB; Validation: SRB and LSW; Writing: SRB, LSW, and BW, with feedback from all other authors; Funding acquisition: BW; Supervision: BW.

## Supplemental Figures

**Figure S1:**
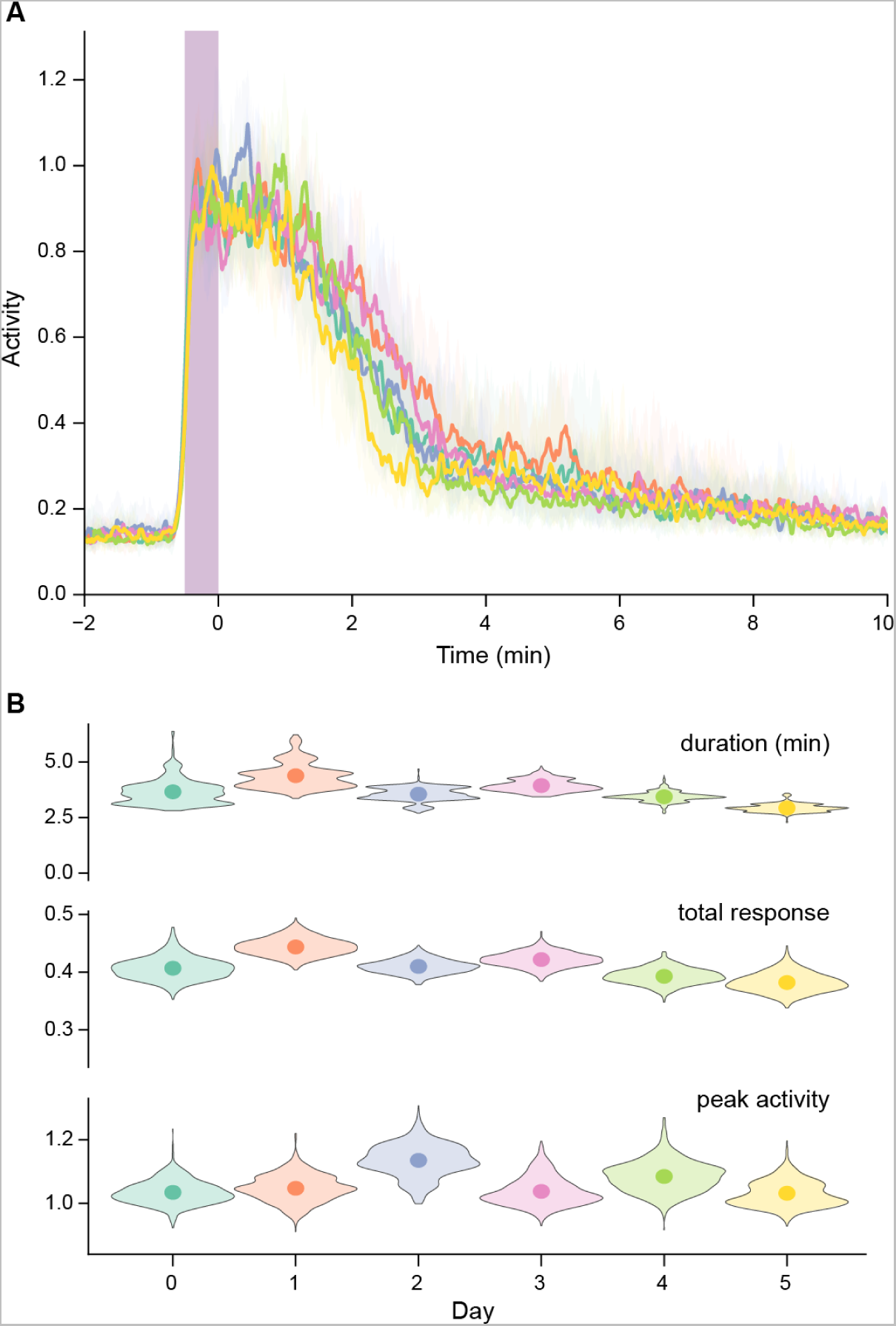
Planarians exhibit consistent behaviors throughout continuous imaging. Related to Figure 1. **(A)** Activity response of whole animals exposed to 30 s UV pulses every 2 hr across days in the imaging chamber. Purple bar: period of UV stimulation. Lines: median activity; shaded region: 99% CI as estimated by 1,000 non-parametric bootstrap samples. **(B)** Quantification of responses shows no change across days. Symbols: mean estimate of measurement on each day; histogram: sampling distribution from 1,000 non-parametric bootstrap samples. No significant differences in the response measurements were found across days relative to the first day in the chamber as determined by 1,000 paired bootstrap samples from each condition (p < 0.01).

**Figure S2:**
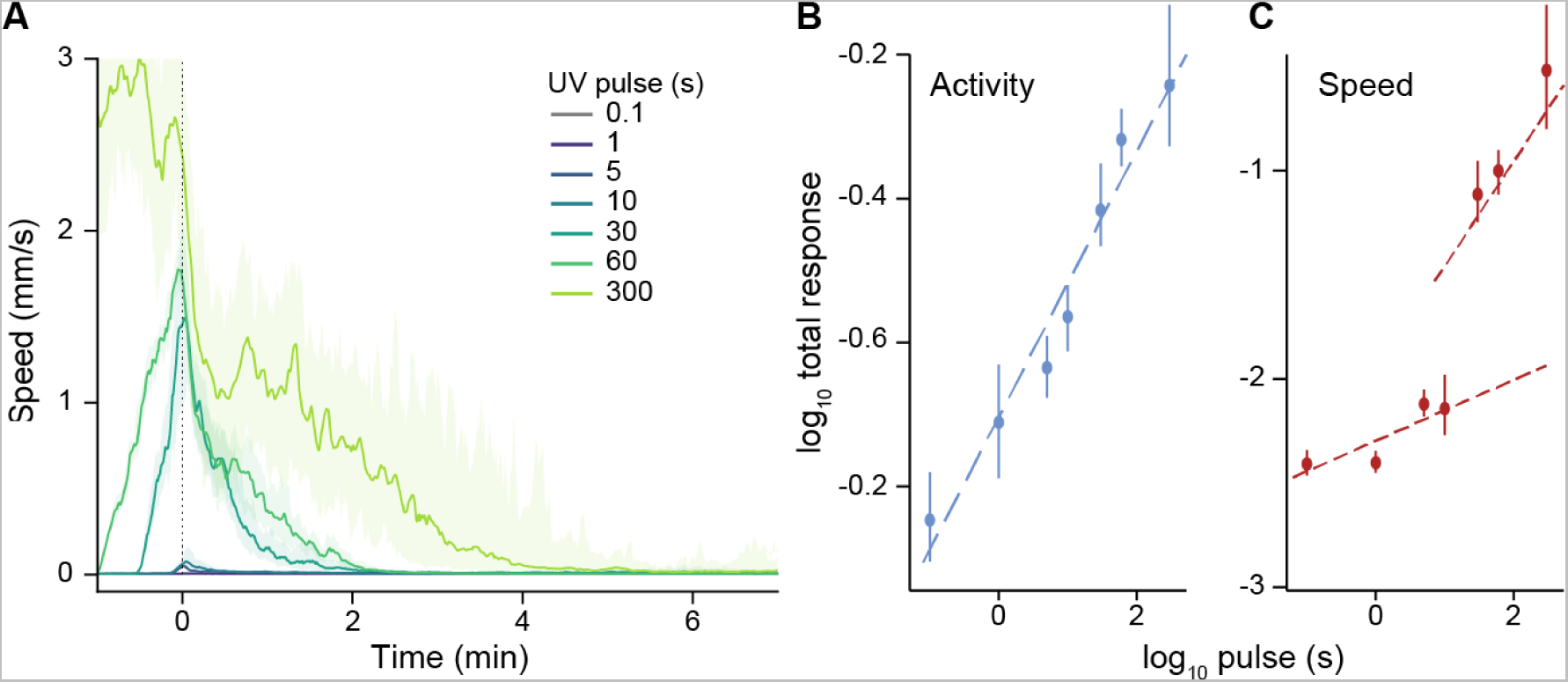
Activity scales as power law with total stimulation. Related to Figure 1. **(A)** Response of whole animals to UV as measured by speed. Time zero: end of UV stimulation. Solid lines: median speed; shaded region 99% CI as estimated by 1000 non-parametric bootstrap samples. **(B)** Total post-stimulus activity follows a power-law scaling with the duration of UV stimulus (slope = 0.18). Error bars: 99% CI. **(C)** Total post-stimulus speed response scales with duration of the UV pulse only in the high-stimulus regime. Error bars: 99% CI.

**Figure S3:**
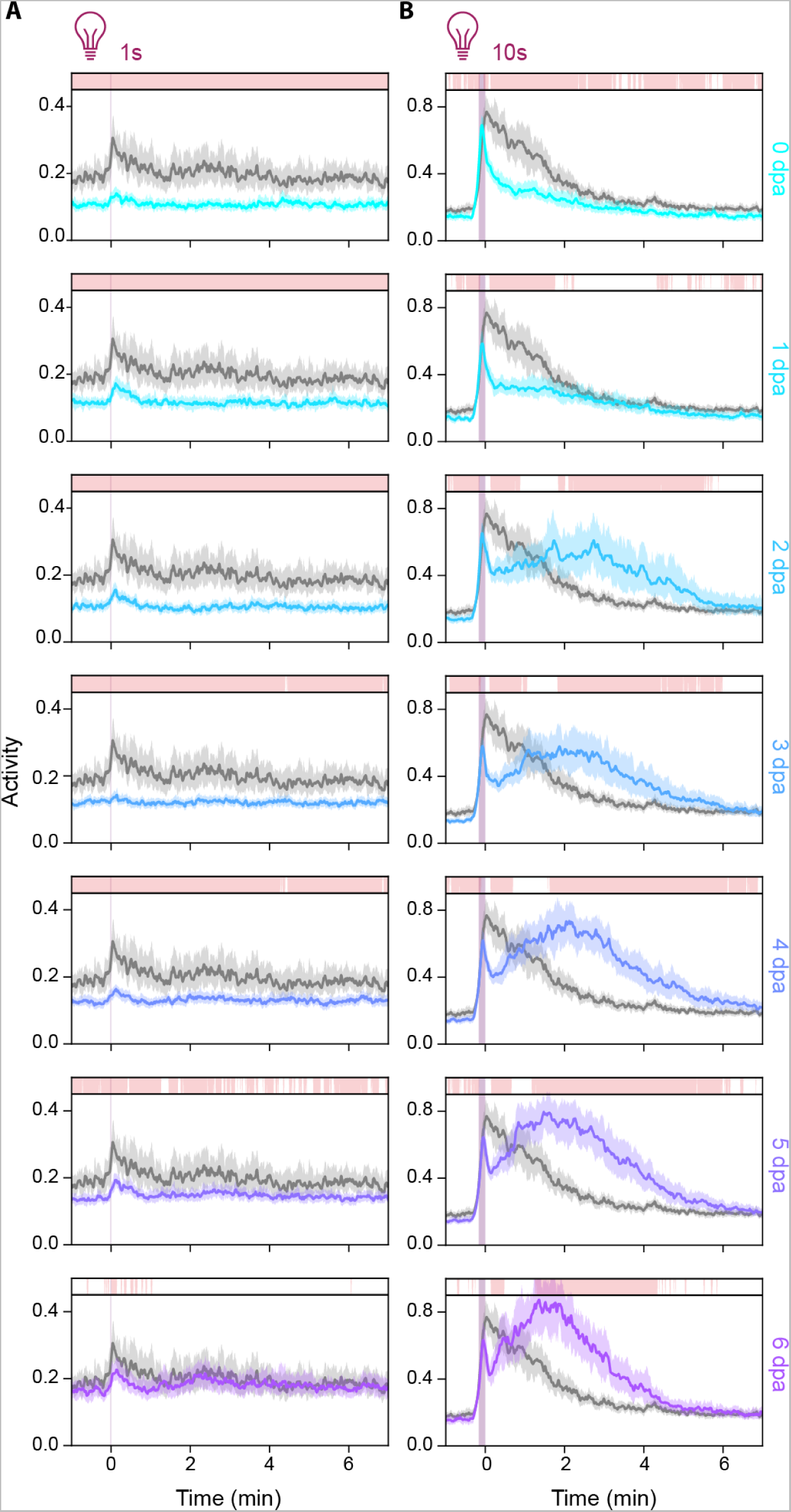
Response to additional UV stimuli after head amputation. Related to Figure 1. Response to 1 s (left) and 10 s (right) UV pulses, by tail fragments after amputation. Gray: whole-animal controls; colored: regenerating tail fragments on each day post amputation (dpa). Bars above the traces indicate time when the response of regenerating animals is significantly different from whole-animal controls as measured by 1,000 nonparametric bootstrap comparisons of the two populations (p < 0.01). Solid lines: median activity; shaded region: 99% CI.

**Figure S4:**
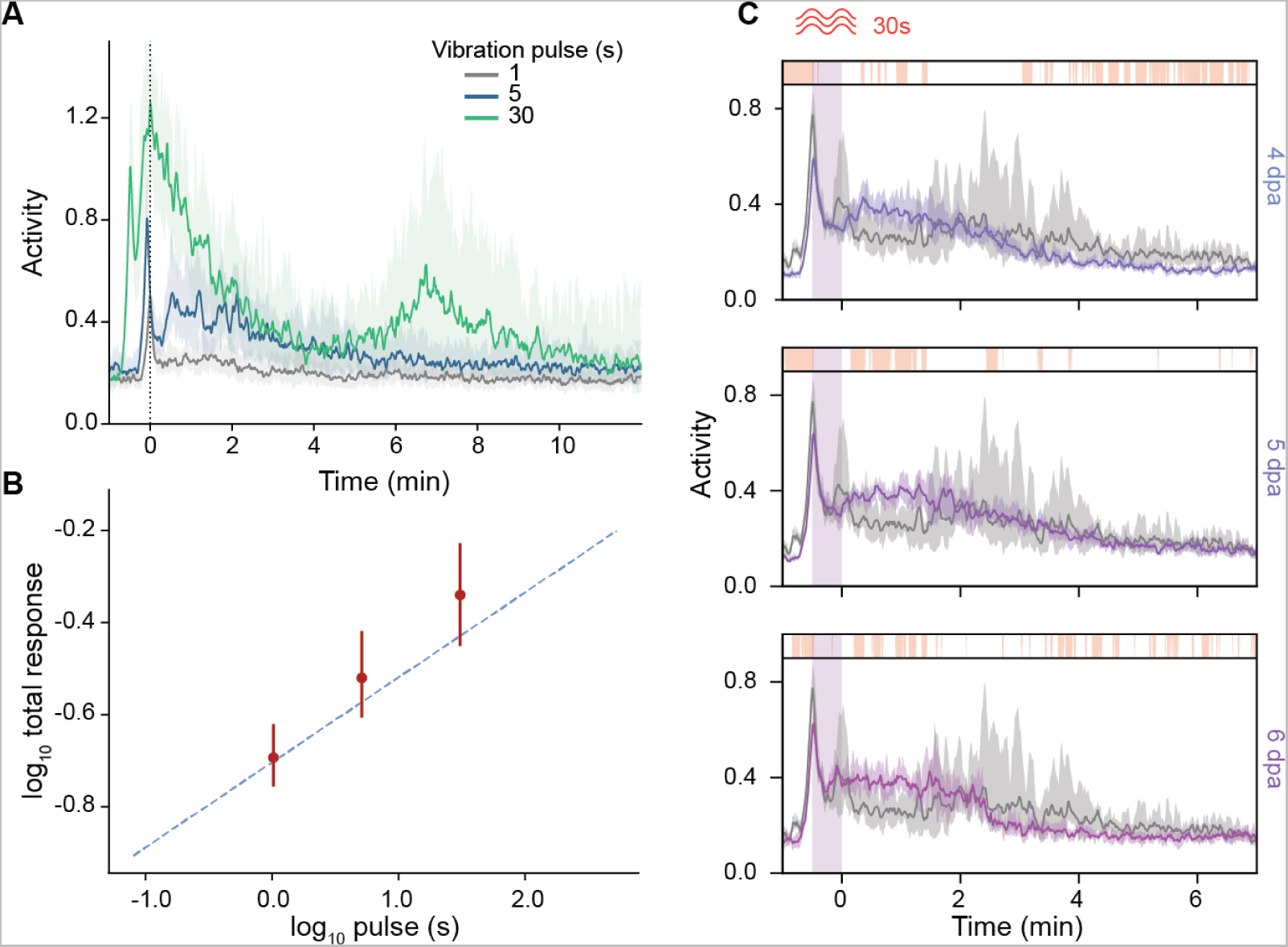
Response to vibration follows similar trends as those to UV. Related to Figure 1. **(A)** Response of whole animals to vibrational pulses. Time zero: end of stimulation. Solid lines: median speed; shaded region 99% CI as estimated by 1,000 non-parametric bootstrap samples. **(B)** Total activity scale with the duration of the vibrational stimulation. Error bars: 99% CI. Dashed line: a power-law scaling with a slope of 0.18 as seen with UV stimulation. **(C)** Response to 30 s vibration in regenerating tail fragments after amputation. Gray: whole-animal response; colored: regenerating response on a given day post amputation (dpa). Bars above the traces indicate time when the response of regenerating animals is significantly different from whole-animal controls as measured by 1,000 nonparametric bootstrap comparisons of the two populations (p < 0.01). For all time traces, lines: median activity; shaded region 99% CI. All confidence intervals are determined from 1000 non-parametric bootstrap samples.

**Figure S5:**
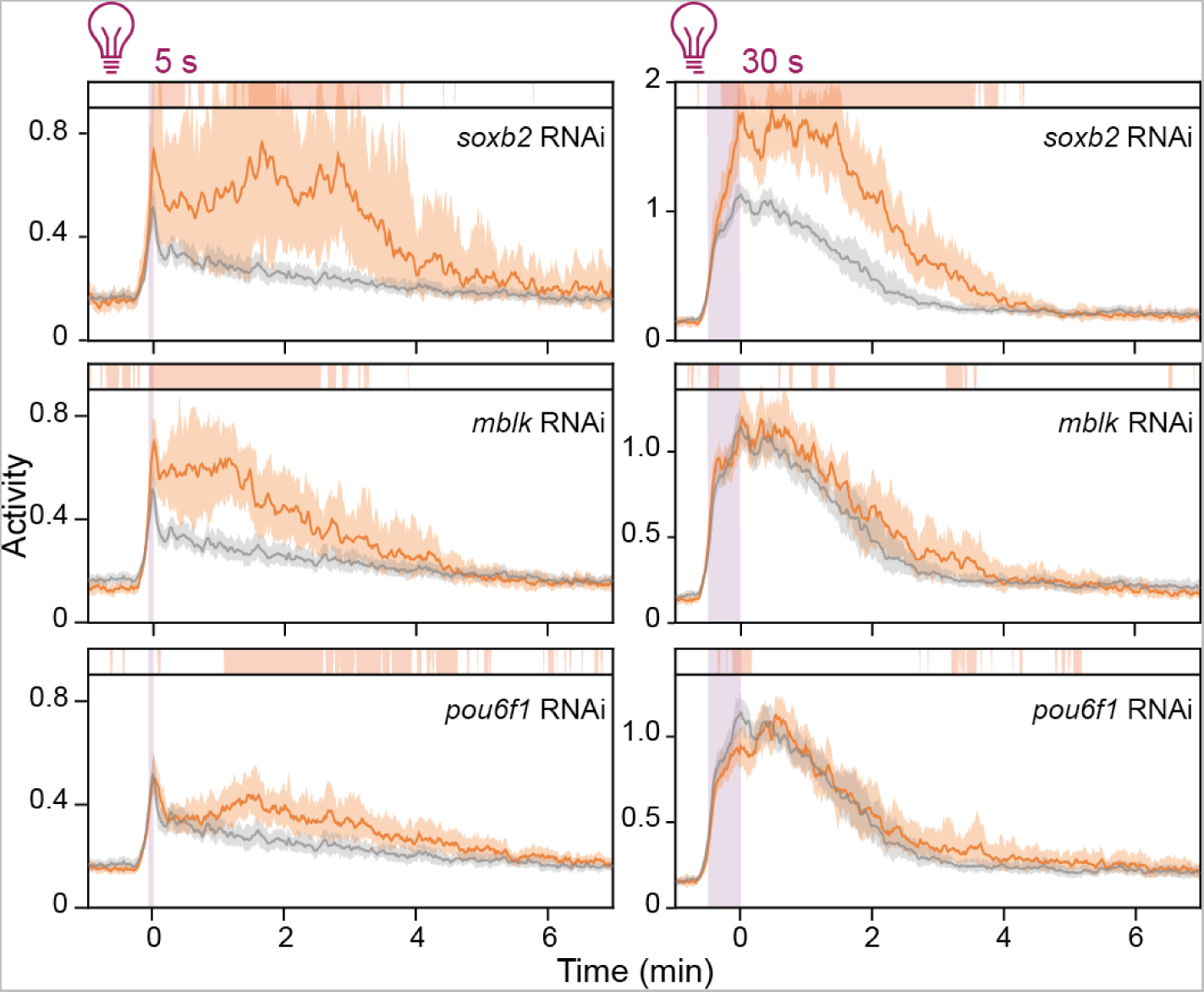
Additional behavioral phenotypes after TF knockdown. Related to Figure 4. Whole-animal response to 5 s (left) and 30 s (right) UV pulses after knockdown of neural TFs. Orange: RNAi conditions; gray: control animals. Bars above the traces indicate time when the response of regenerating animals is significantly different from whole-animal controls as measured by 1,000 nonparametric bootstrap comparisons of the two populations (p < 0.01). Solid lines: median activity; shaded region: 99% CI.

**Figure S6:**
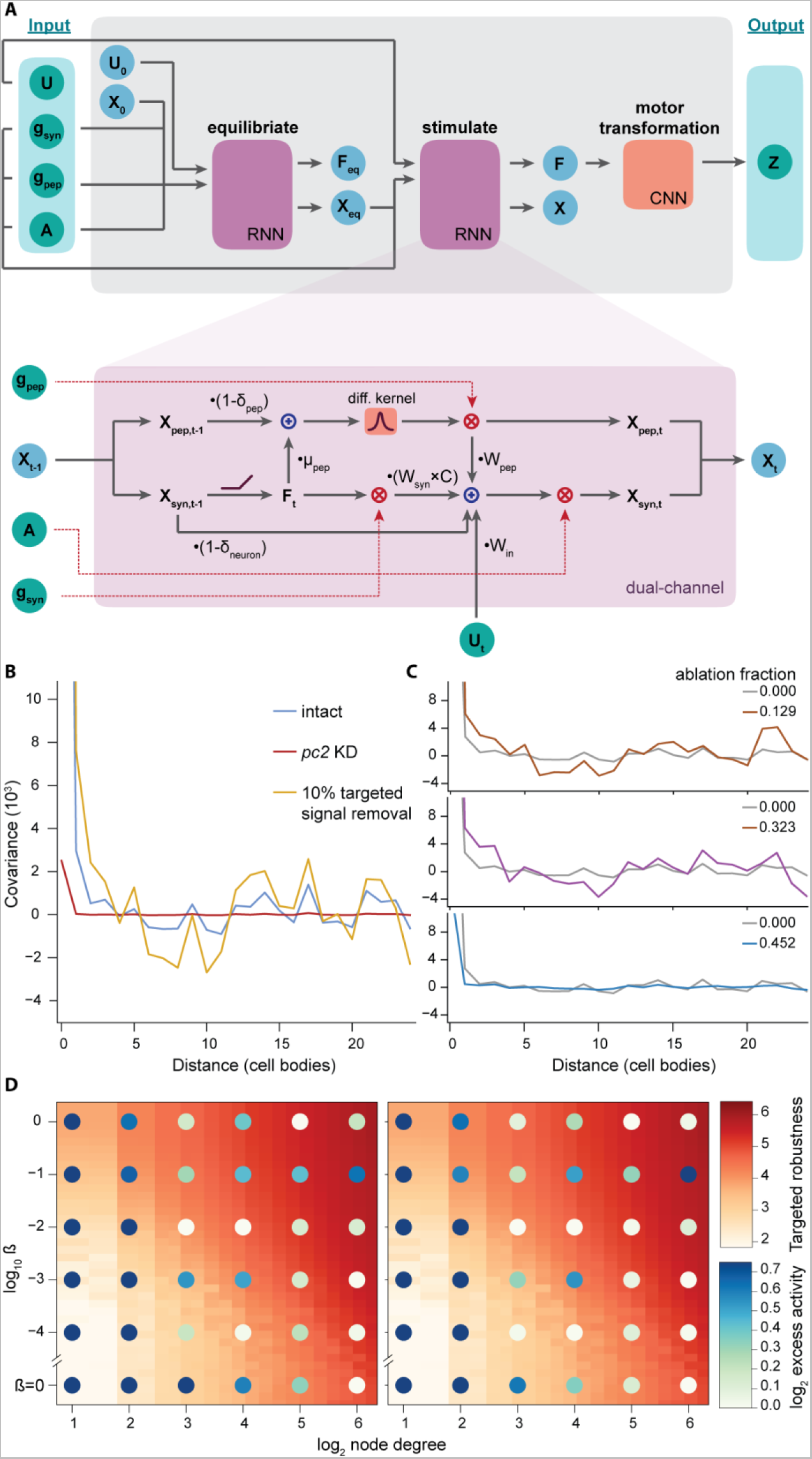
A modified RNN model to take into account the function of neuropeptides. Related to Figure 5. **(A)** Diagram of the dual-channel neuron model. Top: Complete architecture of the model. Bottom: Operations performed at each timestep within the dual-channel RNN cell. **(B)** Spatial covariance of firing rates is driven by peptidergic signals but interfered by small-molecule signals. The initial peak and trough in covariance within the intact model roughly correspond to the diffusive length scale of the excitatory (~3 cell bodies) and inhibitory (~7 cell bodies) peptides, respectively. Removing targeted links increases covariance (gold). Conversely, removing neuropeptides eliminates all spatial covariance (red). **(C)** The magnitude of spatial covariance increases and then decays with increasing ablation. **(D)** Ablation-induced excess activity is dependent on the robustness of connectivity network. Excess activity is defined as the maximum average total response at any ablation level normalized by the total response in the unabated model. Robustness of the connectivity network is estimated using inverse average edge centrality.

## Supplemental files

**Supplemental Movie 1: Activity score captures broad range of animal movements. Related to Figure 1**

The movie records the response of a planarian to a 5 s UV pulse within a droplet. Activity (cyan) resolves non-traversal movements such as nodding and turning better than speed (gray). Shaded region (purple) indicates period of UV stimulation.

**Supplemental Movie 2: *vglut* knockdown causes uncoordinated motor activity. Related to Figure 2**

The movie records the spontaneous behavior of a planarian after *vglut* RNAi. Animals fail to achieve translational movements and display eccentricities of muscular activities.

**Supplemental Movie 3: *pc2* knockdown animals show range of behavior. Related to Figure 2**

Examples of crawling and turning movements in *pc2* RNAi planarians under 30 min continuous UV stimulation. Each clip is from a different animal within the experiment.

**Supplemental Table 1: Primers used in this study. Related to STAR Methods**

Listed are conventional name, contig number, and forward and reverse primers used in cloning experiments for RNAi knockdown.

**Supplemental Table 2: Trial replicate reporting. Related to Figure 1–4**

Number of pulse trials for each experimental condition.

## References

1. Bullmore, E., and Sporns, O. (2009). Complex brain networks: graph theoretical analysis of structural and functional systems. Nat. Rev. Neurosci. 10, 186–198. 10.1038/nrn2575.

2. Aerts, H., Fias, W., Caeyenberghs, K., and Marinazzo, D. (2016). Brain networks under attack: robustness properties and the impact of lesions. Brain 139, 3063–3083. 10.1093/brain/aww194.

3. Dupre, C., and Yuste, R. (2017). Non-overlapping neural networks in Hydra vulgaris. Curr. Biol. 27, 1085–1097. 10.1016/J.CUB.2017.02.049.

4. Weissbourd, B., Momose, T., Nair, A., Kennedy, A., Hunt, B., and Anderson, D.J. (2021). A genetically tractable jellyfish model for systems and evolutionary neuroscience. Cell 184, 5854–5868.e20. 10.1016/J.CELL.2021.10.021.

5. Yasui, K., Kano, T., Standen, E.M., Aonuma, H., Ijspeert, A.J., and Ishiguro, A. (2019). Decoding the essential interplay between central and peripheral control in adaptive locomotion of amphibious centipedes. Sci. Rep. 9, 1–11. 10.1038/s41598-019-53258-3.

6. Lybrand, Z.R., and Zoran, M.J. (2012). Rapid neural circuit switching mediated by synaptic plasticity during neural morphallactic regeneration. Dev. Neurobiol. 72, 1256–1266. 10.1002/DNEU.20993.

7. Le, D., Sabry, Z., Chandra, A., Kristan, W.B., and Collins, E.M.S. (2021). Planarian fragments behave as whole animals. Curr. Biol. 31, 5111–5117.e4. 10.1016/J.CUB.2021.09.056.

8. Li, N., Daie, K., Svoboda, K., and Druckmann, S. (2016). Robust neuronal dynamics in premotor cortex during motor planning. Nature 532, 459–464. 10.1038/nature17643.

9. Hillary, F.G., and Grafman, J.H. (2017). Injured brains and adaptive networks: The benefits and costs of hyperconnectivity. Trends Cogn. Sci. 21, 385–401. 10.1016/J.TICS.2017.03.003.

10. Miranda-Dominguez, O., Mills, B.D., Grayson, D., Woodall, A., Grant, K.A., Kroenke, C.D., and Fair, D.A. (2014). Bridging the gap between the human and macaque connectome: A quantitative comparison of global interspecies structure-function relationships and network topology. J. Neurosci. 34, 5552–5563. 10.1523/JNEUROSCI.4229-13.2014.

11. Niven, J.E., and Laughlin, S.B. (2008). Energy limitation as a selective pressure on the evolution of sensory systems. J. Exp. Biol. 211, 1792–1804. 10.1242/JEB.017574.

12. Nusbaum, M.P., Blitz, D.M., and Marder, E. (2017). Functional consequences of neuropeptide and small-moleculeco-transmission. Nat. Rev. Neurosci. 18, 389. 10.1038/NRN.2017.56.

13. Hille, B. (1992). G protein-coupled mechanisms and nervous signaling. Neuron 9, 187–195. 10.1016/0896-6273(92)90158-A.

14. Fuxe, K., Dahlström, A.B., Jonsson, G., Marcellino, D., Guescini, M., Dam, M., Manger, P., and Agnati, L. (2010). The discovery of central monoamine neurons gave volume transmission to the wired brain. Prog. Neurobiol. 90, 82–100. 10.1016/J.PNEUROBIO.2009.10.012.

15. Flavell, S.W., Pokala, N., Macosko, E.Z., Albrecht, D.R., Larsch, J., and Bargmann, C.I. (2013). Serotonin and the neuropeptide PDF initiate and extend opposing behavioral states in *C. elegans*. Cell 154, 1023–1035. 10.1016/J.CELL.2013.08.001.

16. Brewer, J.C., Olson, A.C., Collins, K.M., and Koelle, M.R. (2019). Serotonin and neuropeptides are both released by the HSN command neuron to initiate *Caenorhabditis elegans* egg laying. PLoS Genet. 15, e1007896. 10.1371/JOURNAL.PGEN.1007896.

17. Van den Pol, A.N. (2012). Neuropeptide transmission in brain circuits. Neuron 76, 98. 10.1016/J.NEURON.2012.09.014.

18. Cebrià, F. (2007). Regenerating the central nervous system: How easy for planarians! Dev. Genes Evol. 217, 733–748. 10.1007/S00427-007-0188-6/FIGURES/6.

19. Evans, D.J., Owlarn, S., Romero, B.T., Chen, C., and Aboobaker, A.A. (2011). Combining classical and molecular approaches elaborates on the complexity of mechanisms underpinning anterior regeneration. PLoS One 6, e27927. 10.1371/JOURNAL.PONE.0027927.

20. Brown, D.D.R., Molinaro, A.M., and Pearson, B.J. (2018). The planarian TCF/LEF factor Smed-tcf1 is required for the regeneration of dorsal-lateral neuronal subtypes. Dev. Biol. 433, 374–383. 10.1016/j.ydbio.2017.08.024.

21. Roberts-Galbraith, R.H., Brubacher, J.L., and Newmark, P.A. (2016). A functional genomics screen in planarians reveals regulators of whole-brain regeneration. Elife 5, e17002. 10.7554/eLife.17002.

22. Currie, K.W., Molinaro, A.M., and Pearson, B.J. (2016). Neuronal sources of hedgehog modulate neurogenesis in the adult planarian brain. Elife 5, e19735. 10.7554/eLife.19735.

23. Vila-Farré, M., and Rink, J.C. (2018). The ecology of freshwater planarians. Methods Mol. Biol. 1774, 173–205. 10.1007/978-1-4939-7802-1_3.

24. Wyss, L.S., Bray, S.R., and Wang, B. (2022). Cellular diversity and developmental hierarchy in the planarian nervous system. Curr. Opin. Genet. Dev. 76, 101960. 10.1016/j.gde.2022.101960.

25. Ross, K.G., Currie, K.W., Pearson, B.J., and Zayas, R.M. (2017). Nervous system development and regeneration in freshwater planarians. Wiley Interdiscip. Rev. Dev. Biol. 6, e266. 10.1002/WDEV.266.

26. Inoue, T., Hoshino, H., Yamashita, T., Shimoyama, S., and Agata, K. (2015). Planarian shows decision-making behavior in response to multiple stimuli by integrative brain function. Zool. Lett. 1, 7. 10.1186/s40851-014-0010-z.

27. Shettigar, N., Chakravarthy, A., Umashankar, S., Lakshmanan, V., Palakodeti, D., and Gulyani, A. (2021). Discovery of a body-wide photosensory array that matures in an adult-like animal and mediates eye-brain-independent movement and arousal. Proc. Natl. Acad. Sci. 118, e2021426118. 10.1073/PNAS.2021426118/SUPPL_FILE/PNAS.2021426118.SM05.MOV.

28. Ross, K.G., Molinaro, A.M., Romero, C., Dockter, B., Cable, K.L., Gonzalez, K., Zhang, S., Collins, E.M.S., Pearson, B.J., and Zayas, R.M. (2018). SoxB1 activity regulates sensory neuron regeneration, maintenance, and function in planarians. Dev. Cell 47, 331–347.e5. 10.1016/J.DEVCEL.2018.10.014.

29. Akiyama, Y., Agata, K., and Inoue, T. (2015). Spontaneous behaviors and wall-curvature lead to apparent wall preference in planarian. PLoS One 10, e0142214. 10.1371/JOURNAL.PONE.0142214.

30. Soitu, C., Feuerborn, A., Tan, A.N., Walker, H., Walsh, P.A., Castrejón-Pita, A.A., Cook, P.R., and Walsh, E.J. (2018). Microfluidic chambers using fluid walls for cell biology. Proc. Natl. Acad. Sci. U. S. A. 115, E5926–E5933. 10.1073/PNAS.1805449115/SUPPL_FILE/PNAS.1805449115.SM05.MP4.

31. Azuma, K., Iwasaki, N., and Ohtsu, K. (1999). Absorption spectra of planarian visual pigments and two states of the metarhodopsin intermediates. Photochem. Photobiol. 69, 99–104. 10.1111/J.1751-1097.1999.TB05312.X.

32. Dexter, J.P., Tamme, M.B., Lind, C.H., and Collins, E.M.S. (2014). On-chip immobilization of planarians for in vivo imaging. Sci. Rep. 4, 6388. 10.1038/SREP06388.

33. Shettigar, N., Joshi, A., Dalmeida, R., Gopalkrishna, R., Chakravarthy, A., Patnaik, S., Mathew, M., Palakodeti, D., and Gulyani, A. (2017). Hierarchies in light sensing and dynamic interactions between ocular and extraocular sensory networks in a flatworm. Sci. Adv. 3, e1603025. 10.1126/SCIADV.1603025.

34. Stevens, S.S. (1957). On the psychophysical law. Psychol. Rev. 64, 153–181.

35. Südhof, T.C. (2013). Neurotransmitter release: The last millisecond in the life of a synaptic vesicle. Neuron 80, 675–690. 10.1016/J.NEURON.2013.10.022.

36. Collins, J.J., Hou, X., Romanova, E. V., Lambrus, B.G., Miller, C.M., Saberi, A., Sweedler, J. V., and Newmark, P.A. (2010). Genome-Wide analyses reveal a role for peptide hormones in planarian germline development. PLoS Biol. 8, e1000509. 10.1371/JOURNAL.PBIO.1000509.

37. Khariton, M., Kong, X., Qin, J., and Wang, B. (2020). Chromatic neuronal jamming in a primitive brain. Nat. Phys. 16, 553–557. 10.1038/s41567-020-0809-9.

38. Jékely, G., Melzer, S., Beets, I., Kadow, I.C.G., Koene, J., Haddad, S., and Holden-Dye, L. (2018). The long and the short of it - A perspective on peptidergic regulation of circuits and behaviour. J. Exp. Biol. 221, jeb166710. 10.1242/JEB.166710/20346.

39. Williams, E.A., Verasztó, C., Jasek, S., Conzelmann, M., Shahidi, R., Bauknecht, P., Mirabeau, O., and Jékely, G. (2017). Synaptic and peptidergic connectome of a neurosecretory center in the annelid brain. Elife 6, e26349. 10.7554/ELIFE.26349.

40. Zhang, X., Pan, H., Peng, B., Steiner, D.F., Pintar, J.E., and Fricker, L.D. (2010). Neuropeptidomic analysis establishes a major role for prohormone convertase-2 in neuropeptide biosynthesis. J. Neurochem. 112, 1168. 10.1111/J.1471-4159.2009.06530.X.

41. Reddien, P.W., Bermange, A.L., Murfitt, K.J., Jennings, J.R., and Sánchez Alvarado, A. (2005). Identification of genes needed for regeneration, stem cell function, and tissue homeostasis by systematic gene perturbation in planaria. Dev. Cell 8, 635–649. 10.1016/J.DEVCEL.2005.02.014.

42. Liu, M., Sharma, A.K., Shaevitz, J.W., and Leifer, A.M. (2018). Temporal processing and context dependency in caenorhabditis elegans response to mechanosensation. Elife 7. 10.7554/ELIFE.36419.

43. Scimone, M.L., Kravarik, K.M., Lapan, S.W., and Reddien, P.W. (2014). Neoblast specialization in regeneration of the planarian Schmidtea mediterranea. Stem Cell Rep. 3, 339–352. 10.1016/J.STEMCR.2014.06.001.

44. Song, H.F., Yang, G.R., and Wang, X.J. (2016). Training excitatory-inhibitory recurrent neural networks for cognitive tasks: A simple and flexible framework. PLoS Comput. Biol. 12, e1004792. 10.1371/JOURNAL.PCBI.1004792.

45. Vyas, S., Golub, M.D., Sussillo, D., and Shenoy, K. V. (2020). Computation through neural population dynamics. Annu. Rev. Neurosci. 43, 249–275. 10.1146/ANNUREV-NEURO-092619-094115.

46. Yang, G.R., Joglekar, M.R., Song, H.F., Newsome, W.T., and Wang, X.J. (2019). Task representations in neural networks trained to perform many cognitive tasks. Nat. Neurosci. 22, 297–306. 10.1038/s41593-018-0310-2.

47. Kapp, F.G., Perlin, J.R., Hagedorn, E.J., Gansner, J.M., Schwarz, D.E., O’connell, L.A., Johnson, N.S., Amemiya, C., Fisher, D.E., Wölfle, U., et al. (2018). Protection from UV light is an evolutionarily conserved feature of the haematopoietic niche. Nature 558, 445–448. 10.1038/s41586-018-0213-0.

48. Karpenko, S., Wolf, S., Lafaye, J., Le Goc, G., Panier, T., Bormuth, V., Candelier, R., and Debrégeas, G. (2020). From behavior to circuit modeling of light-seeking navigation in zebrafish larvae. Elife 9, e52882. 10.7554/ELIFE.52882.

49. Humberg, T.H., Bruegger, P., Afonso, B., Zlatic, M., Truman, J.W., Gershow, M., Samuel, A., and Sprecher, S.G. (2018). Dedicated photoreceptor pathways in *Drosophila* larvae mediate navigation by processing either spatial or temporal cues. Nat. Commun. 9, 1260. 10.1038/s41467-018-03520-5.

50. Lim, M.A., Chitturi, J., Laskova, V., Meng, J., Findeis, D., Wiekenberg, A., Mulcahy, B., Luo, L., Li, Y., Lu, Y., et al. (2016). Neuroendocrine modulation sustains the *C. elegans* forward motor state. Elife 5, e19887. 10.7554/ELIFE.19887.

51. Ma, S., Hangya, B., Leonard, C.S., Wisden, W., and Gundlach, A.L. (2018). Dual-transmitter systems regulating arousal, attention, learning and memory. Neurosci. Biobehav. Rev. 85, 21–33. 10.1016/J.NEUBIOREV.2017.07.009.

52. López-Cruz, A., Sordillo, A., Pokala, N., Liu, Q., McGrath, P.T., and Bargmann, C.I. (2019). Parallel multimodal circuits control an innate foraging behavior. Neuron 102, 407–419.e8. 10.1016/J.NEURON.2019.01.053.

53. Sehgal, A., and Mignot, E. (2011). Genetics of sleep and sleep disorders. Cell 146, 194–207. 10.1016/J.CELL.2011.07.004.

54. Taylor, S.R., Santpere, G., Weinreb, A., Barrett, A., Reilly, M.B., Xu, C., Varol, E., Oikonomou, P., Glenwinkel, L., McWhirter, R., et al. (2021). Molecular topography of an entire nervous system. Cell 184, 4329–4347.e23. 10.1016/J.CELL.2021.06.023.

55. Guillaumin, M.C.C., and Burdakov, D. (2021). Neuropeptides as primary mediators of brain circuit connectivity. Front. Neurosci. 15, 229. 10.3389/FNINS.2021.644313/BIBTEX.

56. Abadi, M., Barham, P., Chen, J., Chen, Z., Davis, A., Dean, J., Devin, M., Ghemawat, S., Irving, G., Isard, M., et al. (2016). TensorFlow: A system for large-scale machine learning. In Proceedings of the 12th USENIX Symposium on Operating Systems Design and Implementation, pp. 265–278.

57. Watts, D.J., and Strogatz, S.H. (1998). Collective dynamics of ‘small-world’ networks. Nature 393, 440–442. 10.1038/30918.

58. Bentley, B., Branicky, R., Barnes, C.L., Chew, Y.L., Yemini, E., Bullmore, E.T., Vértes, P.E., and Schafer, W.R. (2016). The multilayer connectome of Caenorhabditis elegans. PLoS Comput. Biol. 12, 1005283. 10.1371/journal.pcbi.1005283.

59. Hagberg, Aric A; Schult, Daniel A; Swart, P.J. (2008). Exploring network structure, dynamics, and function using NetworkX. In Proceedings of the Python in Science Conference (SciPy), pp. 11–15.

60. Kingma, D.P., and Ba, J. (2015). ADAM: A method for stochastic optimization. In 3rd International Conference for Learning Representations.

61. Werner, S., Rink, J.C., Riedel-Kruse, I.H., and Friedrich, B.M. (2014). Shape mode analysis exposes movement patterns in biology: Flagella and flatworms as case studies. PLoS One 9, e113083. 10.1371/JOURNAL.PONE.0113083.

62. Guan, J., Wang, B., and Granick, S. (2014). Even hard-sphere colloidal suspensions display Fickian yet non-Gaussian diffusion. ACS Nano 8, 3331–3336. 10.1021/NN405476T/SUPPL_FILE/NN405476T_SI_003.AVI.

63. Bishop, C.M. (2006). Pattern recognition and machine learning springer mathematical notation Ni. In, pp. 635–647.

64. Antin, B., Zoltowski, D., Glaser, J., and Linderman, S. (2020). SSM: Bayesian learning and inference for state space models.

65. Kulesa, A., Krzywinski, M., Blainey, P., and Altman, N. (2015). Sampling distributions and the bootstrap. Nat. Methods 12, 477–478. 10.1038/NMETH.3414.

